# The vascular gene *Apold1* is dispensable for normal development but controls angiogenesis under pathological conditions

**DOI:** 10.1101/2022.12.02.518829

**Authors:** Zheng Fan, Raphaela Ardicoglu, Aashil A. Batavia, Ruslan Rust, Lukas von Ziegler, Rebecca Waag, Jing Zhang, Thibaut Desgeorges, Oliver Sturman, Hairuo Dang, Rebecca Weber, Andreas E. Moor, Martin E. Schwab, Pierre-Luc Germain, Johannes Bohacek, Katrien De Bock

## Abstract

The molecular mechanisms of angiogenesis have been intensely studied, but many genes that control endothelial behavior and fate still need to be described. Here, we characterize the role of *Apold1* (Apolipoprotein L domain containing 1) in angiogenesis *in vivo* and *in vitro*. Single-cell analyses reveal that - across tissues - the expression of *Apold1* is restricted to the vasculature, and that *Apold1* expression in endothelial cells (ECs) is highly sensitive to environmental factors. Using *Apold1^-/-^* mice, we find that *Apold1* is dispensable for development and does not affect postnatal retinal angiogenesis nor alters the vascular network in adult brain and muscle. However, when exposed to ischemic conditions following photothrombotic stroke as well as femoral artery ligation, *Apold1^-/-^ mice* display dramatic impairments in recovery and revascularization. We also find that human tumor endothelial cells express strikingly higher levels of *Apold1,* and that *Apold1* deletion in mice stunts the growth of subcutaneous B16 melanoma tumors, which have smaller and poorly perfused vessels. Mechanistically, *Apold1* is activated in ECs upon growth factor stimulation as well as in hypoxia, and *Apold1* intrinsically controls EC proliferation but not migration. Our data demonstrate that *Apold1* is a key regulator of angiogenesis in pathological settings, whereas it does not affect developmental angiogenesis, thus making it a promising candidate for clinical investigation.

## Introduction

Angiogenesis, the formation of new blood vessels from existing ones, is crucial for normal embryonic development and thus essential for life [1–3]. In healthy tissues, however, angiogenesis only occurs in the muscle [4] and in specific, highly activated brain areas following exercise training [5] and in the uterus where it ensures endometrial growth during the menstrual cycle [6, 7]. Angiogenesis is a strictly timed, multi-step process that is tightly controlled by an intimate interplay between several pro- and anti-angiogenic growth factors that regulate the activity of complex cellular signaling networks [8]. Those ultimately orchestrate changes in the function and fate of endothelial cells (ECs), the main cell type composing the vascular wall. Decades of intense research, mostly using models of developmental angiogenesis, have led to the discovery of a plethora of angiogenic master regulators[2, 3]. But undoubtedly, many genes that control EC behavior, fate, and/or EC interactions with other cells to ensure the generation of functional blood vessels, still need to be described [9].

In many pathological conditions, angiogenesis plays a crucial role. On the one hand, angiogenesis secures tissue viability and recovery during and after injuries often associated with hypoxia, such as in wounds, upon ischemia or after stroke, thereby ensuring tissue repair [10–12]. On the other hand, the "angiogenic switch" in cancer fuels tumor growth and malignancy [13, 14]. In pathological conditions, angiogenesis often relies on the reactivation of developmental pathways that control physiological angiogenesis [15, 16]. For example, many tumors activate the expression of vascular endothelial growth factor (VEGF), a master regulator of physiological angiogenesis, in response to environmental signals such as hypoxia or following oncogenic (or loss of tumor suppressor function) mutations. As a result, anti-angiogenic drugs targeting VEGF, but also other crucial regulators of developmental angiogenesis, have been extensively tested for anti-cancer treatment. Despite some successes, their efficacy to slow down tumor progression and improve patient survival remains limited due to serious side-effects, toxicity and acquired resistance [13, 17–19]. Therefore, identifying novel molecular targets that are redundant for physiological angiogenesis and endothelial survival, but specifically regulate pathological angiogenesis, holds great therapeutic potential.

*Apold1* (Apolipoprotein L domain containing 1, also known as Verge) is a gene that is thought to be selectively expressed in ECs [20]. *Apold1* expression rapidly increases under metabolically demanding conditions, e.g., in the rat heart after vigorous physical activity [21], in human muscle after acute aerobic and resistance exercise [22, 23], and in the rodent brain in response to activating/stressful stimuli [24, 25] and seizures [20]. It appears that this response is driven by increased oxygen demand, as *Apold1* mRNA increases in ischemic brains [20, 26], and under hypoxic conditions in various tissues [27–30]. Angiogenic growth factors such as FGF2 and (to a lesser extent) Angiopoietin-2 also activate *Apold1* [20]. But while *in vitro* studies suggested that *Apold1* controls vascular permeability [20, 31] and the secretion of Weibel Palade bodies [31], little is known about the functional role of *Apold1 in vivo*. In this respect, a mutation in *Apold1* was recently described in a family of patients with a novel inherited bleeding disorder [31], yet *Apold1* knockout mice (*Apold1^-/-^*) present with higher platelet reactivity and a prothrombotic phenotype [32]. In a mouse model of neonatal stroke, *Apold1^-/-^* pups showed reduced angiogenesis after stroke and impaired long-term functional recovery [26]. However, in adult mice, where *Apold1* expression is much lower, acute stroke caused similar size lesions and comparable functional impairment in *Apold1^-/-^* mice and wild-type controls [33]. Importantly, the role of *Apold1* in recovery from stroke, including angiogenesis and revascularization, was not assessed. Thus, whether *Apold1* contributes to angiogenesis in adulthood, is not known.

Here we investigated the role of *Apold1* in angiogenesis *in vivo* and *in vitro*. We report that *Apold1* is dispensable for developmental angiogenesis, but it is necessary for functional revascularization during recovery from ischemia in the central nervous system and in the periphery. We also find that human tumor ECs express strikingly higher levels of *Apold1, and* that *Apold1* deletion in mice reduces tumor growth by limiting EC proliferation.

## Results

### *Apold1* is expressed in endothelial cells and regulated by metabolic demand

*Apold1* was originally reported to be an immediate early gene expressed in endothelial cells (ECs), primarily during embryogenesis [20]. To assess *Apold1* expression in adult animals, we first interrogated "Tabula Muris", a large single-cell RNA sequencing (scRNAseq) compendium covering 20 mouse organs [34]. Across organs, *Apold1* expression was largely restricted to ECs (Fig. S1A-C). When narrowing the analysis to individual, highly vascularized tissues, *Apold1* was highly expressed in ECs in the brain (Fig. S1D), muscle (Fig. S1E) and liver (Fig. S1F). As ECs differ substantially between tissues [35], we took a closer look at *Apold1* mRNA in both brain and muscle, two tissues that differ radically in cellular complexity and metabolic profile. Reanalysis of a scRNAseq dataset from mouse adult skeletal hindlimb muscle [36] confirmed that *Apold1* expression is restricted to vascular cells (Fig. 1A-C), with abundant expression in ECs, but also to a lesser extent in pericytes and smooth muscle cells (Fig. 1B,C). To better understand the distribution of *Apold1* within the different types of ECs within muscle, we evaluated *Apold1* expression in FACS-sorted Pecam1^+^CD45^-^ ECs that were collected from the mouse gastrocnemius muscle [37] (Fig. 1D). *Apold1* was widely distributed across the different EC populations ranging from arterial to capillary and venous ECs (Fig. 1E). Interestingly, its expression in capillary ECs derived from white, glycolytic muscle (WmECs), which have low angiogenic potential, was noticeably lower when compared to capillary ECs derived from red, oxidative muscle (RmECs) with high angiogenic potential (Fig. 1E). To confirm this, we isolated WmECs (from EDL) and RmECs (from Soleus) using FACS (Pecam1^+^CD45^-^) and showed with RT-qPCR that *Apold1* was enriched in RmECs compared to WmECs (Fig. 1F). We then tested whether exercise, an intervention known to activate RmECs, would affect *Apold1* expression (Fig. 1G). Following two weeks after daily voluntary wheel running, we again isolated Pecam1^+^CD45^-^ ECs using FACS from calf muscle. RT-qPCR showed a marked increase in *Apold1* expression in whole muscle as well as in sorted mECs after exercise (Fig. 1H).

**Fig. 1.**
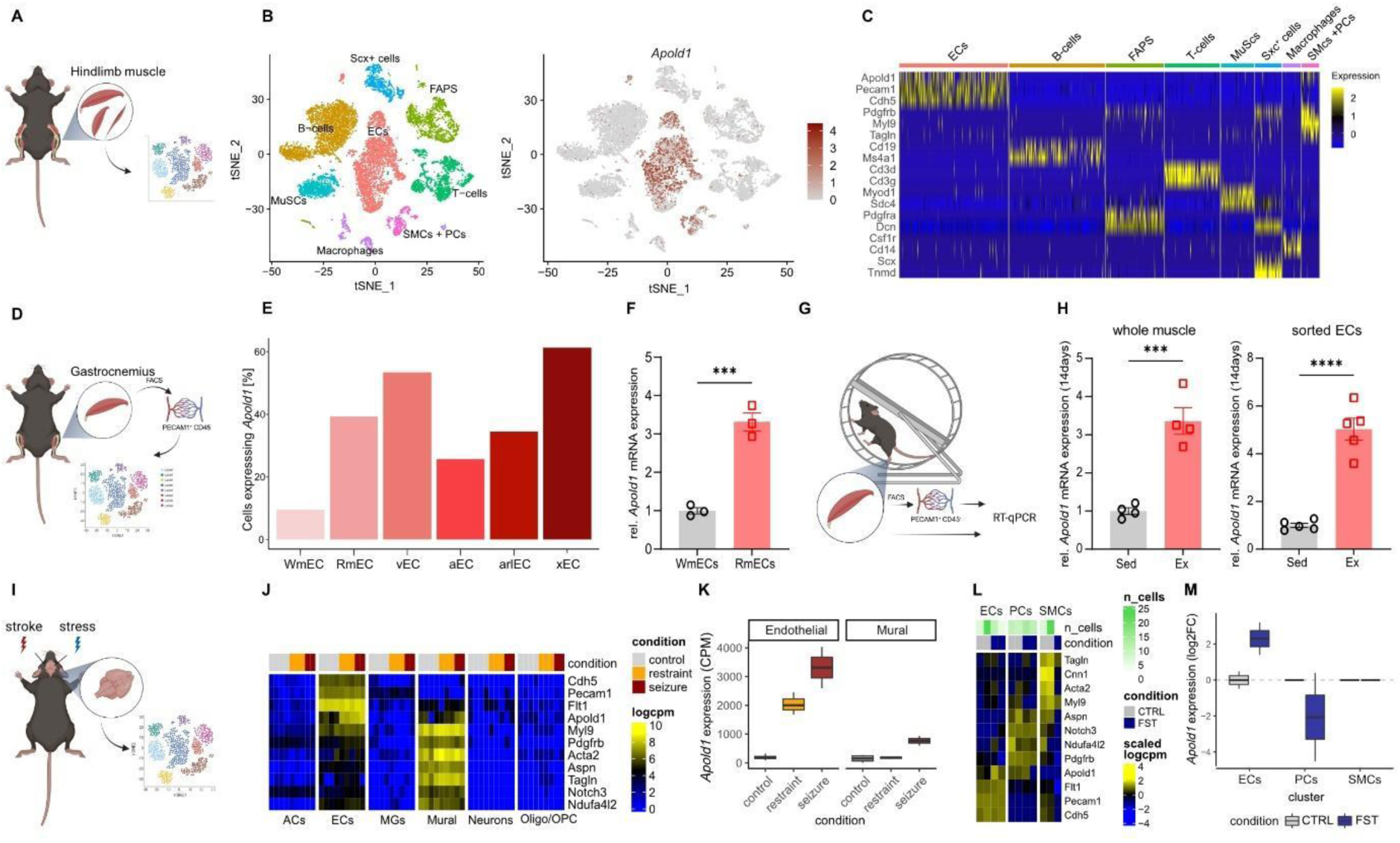
*Apold1* gene expression in muscle and brain is restricted to the vasculature, including endothelial cells, pericytes and smooth muscle cells and triggered by environmental stimuli. (A) Experimental design [36]. (B) t-Stochastic neighbor embedding (tSNE) plots of distribution of main muscle-resident populations reanalyzed [36] (ECs -endothelial cells; PCs - pericytes; SMCs -smooth muscle cells). (C) RNA expression heat map for given cell populations (column) and genes (row) sorted by population specific gene expression reanalyzed [36]. (D) Experimental design [37]. (E) Percentage of *Apold1* RNA expression in all endothelial populations reanalyzed [37]. (F) Relative *Apold1* mRNA expression in WmECs and RmECs (n (WmECs/RmECs*^-^*) = 3/3). (G) Experimental design. (H) Relative *Apold1* mRNA expression in whole muscle and sorted ECs after 14 days of voluntary wheel running (n (Sed/Ex) = 4/4). (I) Experimental design [39, 41]. (J) RNA expression heat map for control, restraint stress and seizure conditions (column) and genes (rows) reanalyzed [39]. (L) RNA expression heat map of ECs, PCs and SMCs in mice brain exposed to control (CTRL) and forced swim test conditions (FST) [41]. (M) *Apold1* RNA expression upon forced swim test (FST) in ECs, PCs and SMCs of the brain compared to control conditions [41]. Student’s t test in F and H (***p< 0.001, ****=p<0.0001). The data shown are mean ± SEM.

We then assessed *Apold1* expression in the brain and turned to an extensive single-cell resource on the mouse nervous system [38], which confirmed that *Apold1* expression is restricted to vascular cells including ECs and to a lesser extent mural cells (pericytes and smooth muscle cells) (Fig. S2). It had previously been reported that metabolically demanding environmental insults like seizures, stress or exercise increase *Apold1* expression in brain tissue [20, 21, 24, 25]. To explore in which cell types this regulation occurs, we used a scRNAseq study that compared mice that had been injected with the seizure-inducing agent pentylenetetrazole (PTZ, which triggers strong pathological levels of neuronal activity), exposed to immobilization stress (which triggers high but physiological levels of neuronal activity), or received no treatment [39] (Fig. 1I). This dataset contained several biological replicates per group, allowing us to quantitatively compare multi-condition scRNAseq datasets [40]. We found that *Apold1* expression was again restricted to vascular cells, including ECs and mural cells (Fig. 1J), and that seizures triggered robust *Apold1* expression mainly in ECs, although we also observed a small increase in mural cells (Fig. 1K). In addition, immobilization stress triggered *Apold1* expression only in ECs but not in mural cells (Fig. 1K), suggesting that *Apold1* expression in ECs is particularly sensitive to environmental stimuli. To further examine whether *Apold1* regulation is indeed more sensitive in ECs than in mural cells, we turned to a single-nucleus RNA sequencing experiment (snRNAseq), in which we had assessed the transcriptomic response of mice to an acute swim stress exposure [41]. Indeed, *Apold1* was expressed in ECs and pericytes, but the stress-induced increase in *Apold1* expression was restricted to ECs (Fig. 1L,M). These analyses demonstrate that *Apold1* expression across the organism is restricted to vascular cells, and that *Apold1* expression is readily stimulated by environmental challenges, particularly in ECs.

### Loss of *Apold1* does not impair normal development

To assess whether *Apold1* plays a role in EC function *in vivo*, we turned to *Apold1* knockout (*Apold1^-/-^*) mice. Because of the key role of angiogenesis during development [42], and because *Apold1* expression is high during development [20], we first assessed whether *Apold1^-/-^* mice display developmental defects. We set up several heterozygous (*Apold1^+/-^*) breeding pairs and observed the expected Mendelian distribution of offspring genotypes with very similar distributions in males and females (Fig. 2A). Also, *Apold1^-/-^* females were able to produce normal numbers of healthy pups when mated with wild-type (WT) or *Apold1^-/-^* males, and we noticed no increased incidence of natural deaths in *Apold1^-/-^* relative to wild-type WT mice (data not shown). Moreover, mice had similar body weight (Fig. 2B). *Apold1^-/-^* mice as well as heterozygous *Apold1^+/-^* mice appeared developmentally normal and were behaviorally indistinguishable from WT littermates in terms of locomotor activity, exploratory behaviors and emotionality in the open field test and the light-dark box test (Fig. 2C). To investigate whether loss of *Apold1* leads to more subtle changes in developmental angiogenesis, we subsequently used the postnatal retina, which allows the characterization of distinct changes in EC fate [43, 44]. At postnatal day 5 (P5), whole-mount isolectin-B4-stained retinas showed that vascular parameters are normal in *Apold1^-/-^* mice, with normal vascular outgrowth, number of branch points, total vessel area and number of tip cells in *Apold1^-/-^* mice when compared to their WT littermates (Fig. 2D-E). Further, the analysis of the vasculature in the oxidative as well as the glycolytic part of the gastrocnemius muscle of adult animals showed no alterations in terms of vessel area (Fig. 2 F-G).

**Figure 2.**
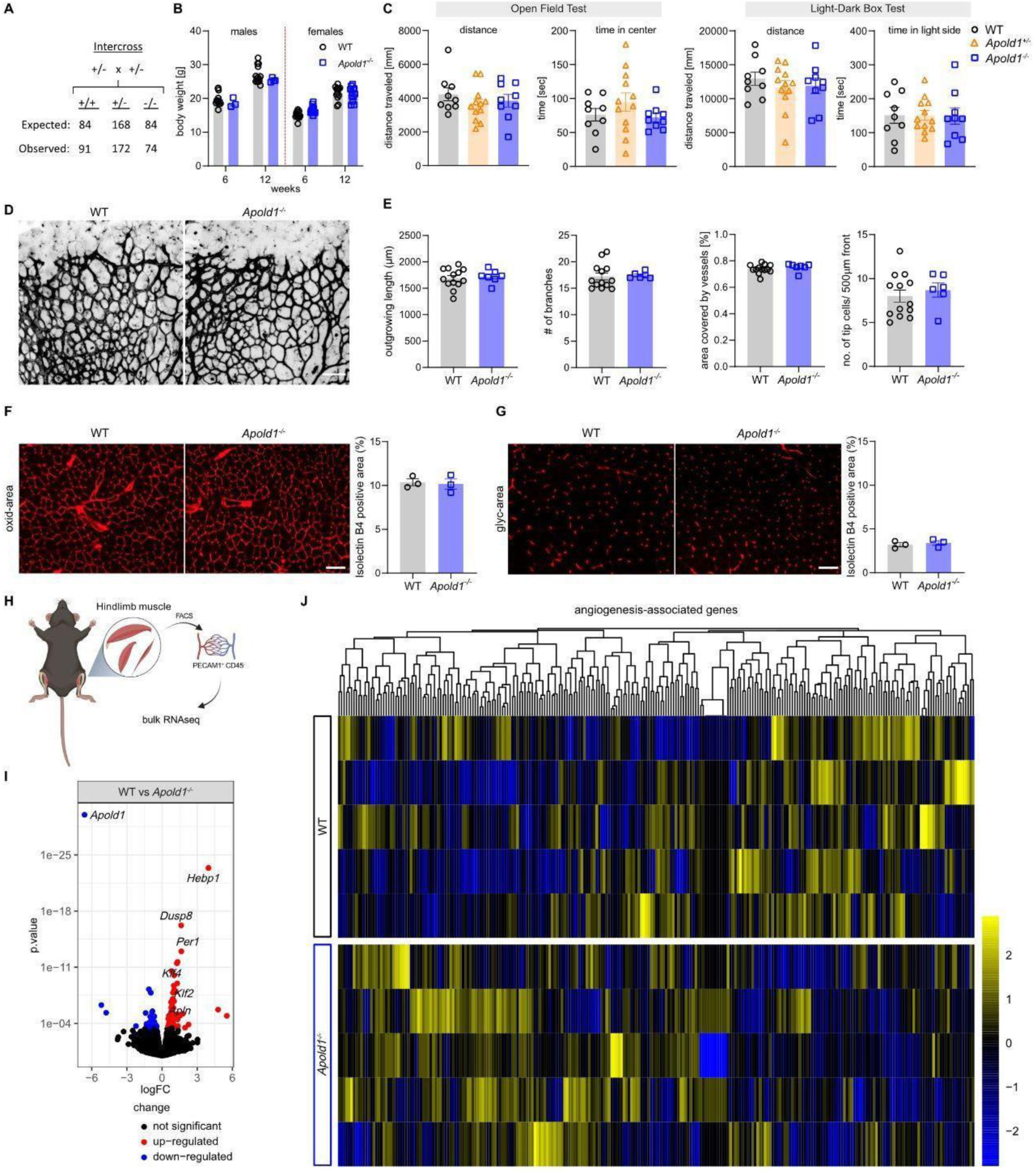
Apold1 deficiency does not affect physiological angiogenesis. (A) Expected and observed Mendelian distribution of heterozygous breedings (Chi-square test: 0.9585; p= 0.6193). (B) Quantification of body weight at 6 and 12 weeks of age in females and males between genotypes (n = 3-14 per group). (C) Behavioral analysis of WT and *Apold1^-/-^* in the open-field test and light-dark box measured by distance traveled and time in center/time in light side (n (WT/*Apold1^+/-^*/*Apold1^-/-^*) = 9/12/9). (D) Representative images of isolectin-B4-stained (black) retinal vessels of postnatal day 5 (P5) pups. Scale bar, 100 µm. (E) Quantification of outgrowing vessel length, number of branches, percentage of area covered by vessels, and number of tip cells in 500 µm front of the retina (n (WT/*Apold1^-/-^*) = 13/6). (F) Representative images and percentage positive area of blood vessels stained with isolectin B4 (red) in the oxidative area of the gastrocnemius (oxid-area) (n (WT/*Apold1^-/-^*) = 3/3). Scale bar, 200 μm. (G) Representative images and percentage of positive area of blood vessels stained with isolectin B4 in the glycolytic area of the gastrocnemius (glyc-area) (n (WT/*Apold1^-/-^*) = 3/3). Scale bar 200 µm. (H) Experimental design. (I) Volcano plot of significantly differentially expressed genes by bulk RNAseq of WT vs *Apold1^-/-^* mECs (n (WT/*Apold1^-/-^*) = 5/5). (J) RNA expression heat map of all angiogenesis-associated genes (n(WT/*Apold1^-/-^*) = 5/5). Student’s test in B, E, F, and G; One-way ANOVA with Tukey’s multiple comparison test in C. The data shown are mean ± SEM.

Screening of the transcriptomic profile by bulk RNAseq of WT and *Apold1^-/-^* isolated muscle ECs revealed minor transcriptional differences, as only 89 genes were differentially expressed (Fig. 2H-I). Amongst those were *Klf2* and *Klf4,* known regulators of vascular integrity by shear stress [45, 46], but activation of their downstream targets was not obvious (Fig. 2I; see list of differentially expressed genes in Supplementary Table1). We also observed an increase in *Apln* (a gene with pleiotropic functions in ECs, reported to promote angiogenesis [47] (Fig. 2I), but we could not detect any concerted regulation of angiogenesis-related genes (Fig. 2J). Altogether, these data show that loss of *Apold1* does not lead to any obvious developmental deficits in overall health, and that *Apold1* is dispensable for developmental angiogenesis.

### Loss of Apold1 impairs recovery from stroke

To subsequently study the role of *Apold1* during angiogenesis in pathological settings, we turned to a mouse model of stroke. We first tested whether photothrombotic (i.e. microthrombotic focal) strokes in the sensorimotor cortex of adult mice would lead to changes in *Apold1* mRNA expression in the ischemic border region, where angiogenesis is induced [48] (Fig. 3A). As expected, *Apold1* expression increased in the microdissected peri-infarct tissue 2 days after stroke and returned to baseline levels within a week (Fig. 3A,B, right). *Apold1* expression did not change in the intact contralateral cortex (Fig. 3A,B, left). Then, we induced photothrombotic strokes in *Apold1^-/-^* mice and WT littermates, using either a weaker or a stronger paradigm to generate milder and more severe strokes (Fig. 3C). Stroke volume was similar in *Apold1^-/-^* and WT mice (Fig. 3E), indicating that loss of *Apold1* does not impact the severity of an ischemic injury, in agreement with previous findings [33]. Also, the non-injured (contralateral) cortex of *Apold1^-/-^* mice had a normal vascular network (vascular area fraction, number of branches and vascular length), consistent with normal vascular development of *Apold1^-/-^* mice (Fig. 3D,F). However, 21 days after the stroke, we observed a reduction in vascular area, branch numbers and total length of the vascular network in the peri-infarct border zone of *Apold1^-/-^* mice as compared to the corresponding region of WT mice (Fig. 3F). Labeling of proliferating cells one week after stroke using 5-ethynyl-2’-deoxyuridine (EdU) injections combined with immunofluorescent labeling of PECAM1^+^ ECs showed that *Apold1^-/-^* ECs within the ischemic border zone proliferate less (Fig. 3G,I). Thus, *Apold1^-/-^* mice show impaired angiogenesis in the ischemic stroke border zone during the three-week recovery period.

**Figure 3.**
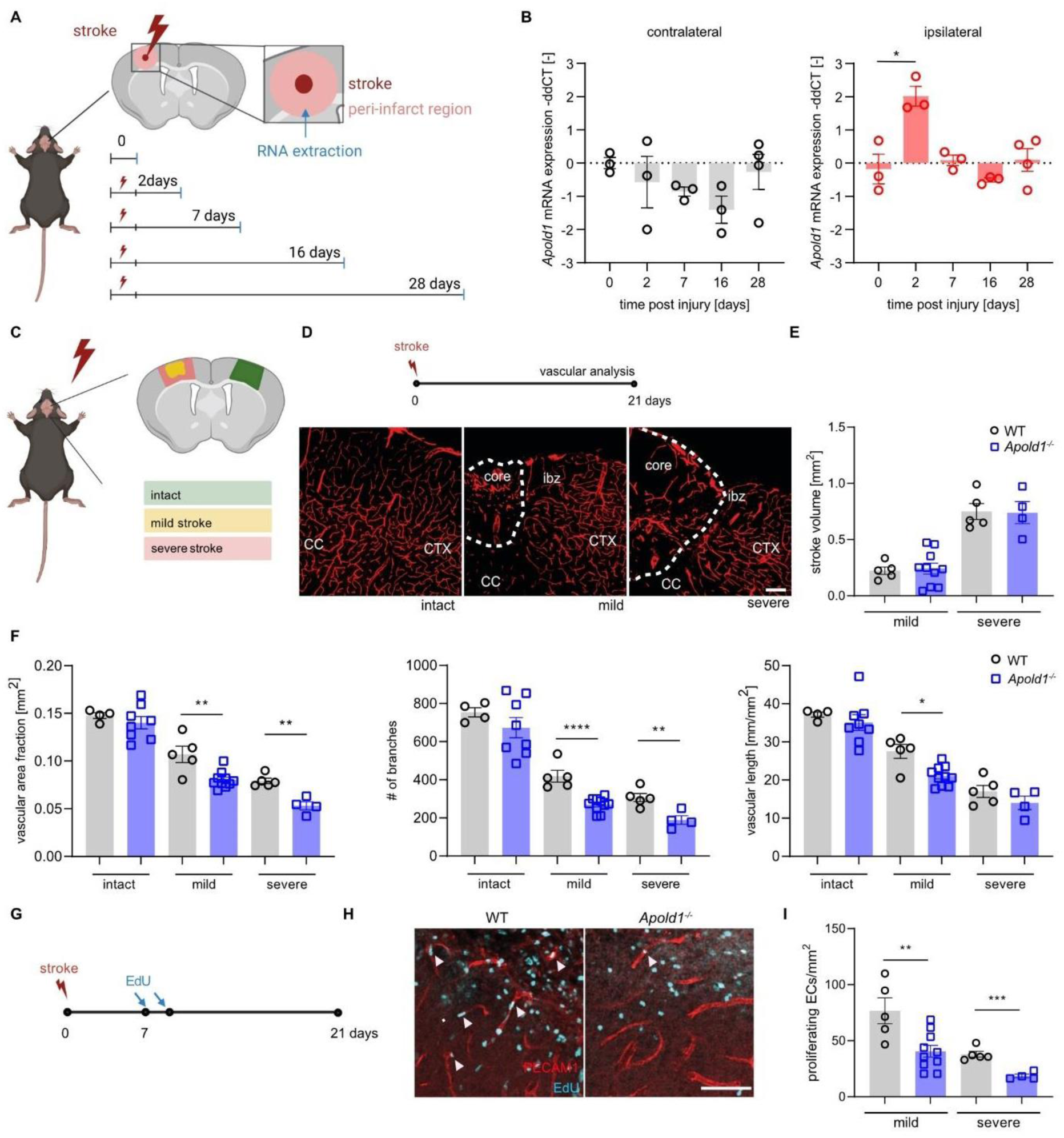
Apold1 is required for angiogenesis and re-vascularization after photothrombotic stroke. (A) Experimental design. (B) Time course of relative *Apold1* mRNA expression on contralateral and ipsilateral cortex (n =3-4 per timepoint). (C) Illustration of stroke size and location. (D) Experimental design and representative images of intact and injured PECAM1^+^ vasculature. (E) Stroke volumes 21 days following injury (n =4-10 per group; Scale bar, 100 µm). (F) Quantitative analysis of vascular density, number of branches, length of blood vessels in the ischemic border regions (n =4-8 per group). (G) Experimental design. (H) Representative images of newly formed vascular cells by PECAM1^+^/EdU^+^ co-staining in WT and *Apold1^-/-^* animals. Scale bar, 50 µm. (I) Quantification of proliferating ECs (n =4-10 per group). Two-way ANOVA with Tukey’s multiple comparisons test in B; Student’s t-test in E, F and G (*p< 0.05; **p< 0.01; ***p<0.001; ****p<0.0001). The data shown are mean ± SEM.

### Apold1 is required for revascularization after hindlimb ischemia

Because the regenerative capacity of peripheral tissues is much greater than that of the central nervous system (CNS), and because *Apold1* is abundantly expressed in ECs from non-CNS vascular beds (see Figs. 1 and S1), we asked whether *Apold1^-/-^* mice also show impaired angiogenesis in the regeneration of peripheral tissue. We used hindlimb-ischemia, where ligation of the femoral artery strongly reduces perfusion of the hindlimb and causes severe hypoxia followed by a robust angiogenic response [49, 50]. Analysis of *Apold1* mRNA showed a strong (8-fold) increase in whole tissue isolated from the ipsilateral (ischemic) muscle compared to the contralateral side 12 hours after femoral artery ligation (Fig. 4A; 4B, left). Isolation of muscle endothelial cells (mECs) from the ligated hindlimb 3 days after induction of ischemia revealed a nearly 20-fold increase in *Apold1* mRNA content when compared to the non-ligated contralateral side (Fig. 4B, right). *In situ* hybridization confirmed a strong increase in *Apold1* expression after ischemia, colocalized with *Pecam1* (Fig. 4D). Laser doppler imaging showed that blood flow was similarly reduced upon femoral artery ligation in both WT and *Apold1^-/-^* mice, but while it gradually recovered in WT mice during the 4-week follow-up period, reperfusion remained severely impaired in *Apold1^-/-^* mice (Fig. 4E,F). This impairment was observed in both sexes, and the deficit was similarly profound in *Apold1^-/-^* and in mice haplo-deficient for *Apold1* (*Apold1^-/+^*) (Fig. 4F).

**Fig. 4.**
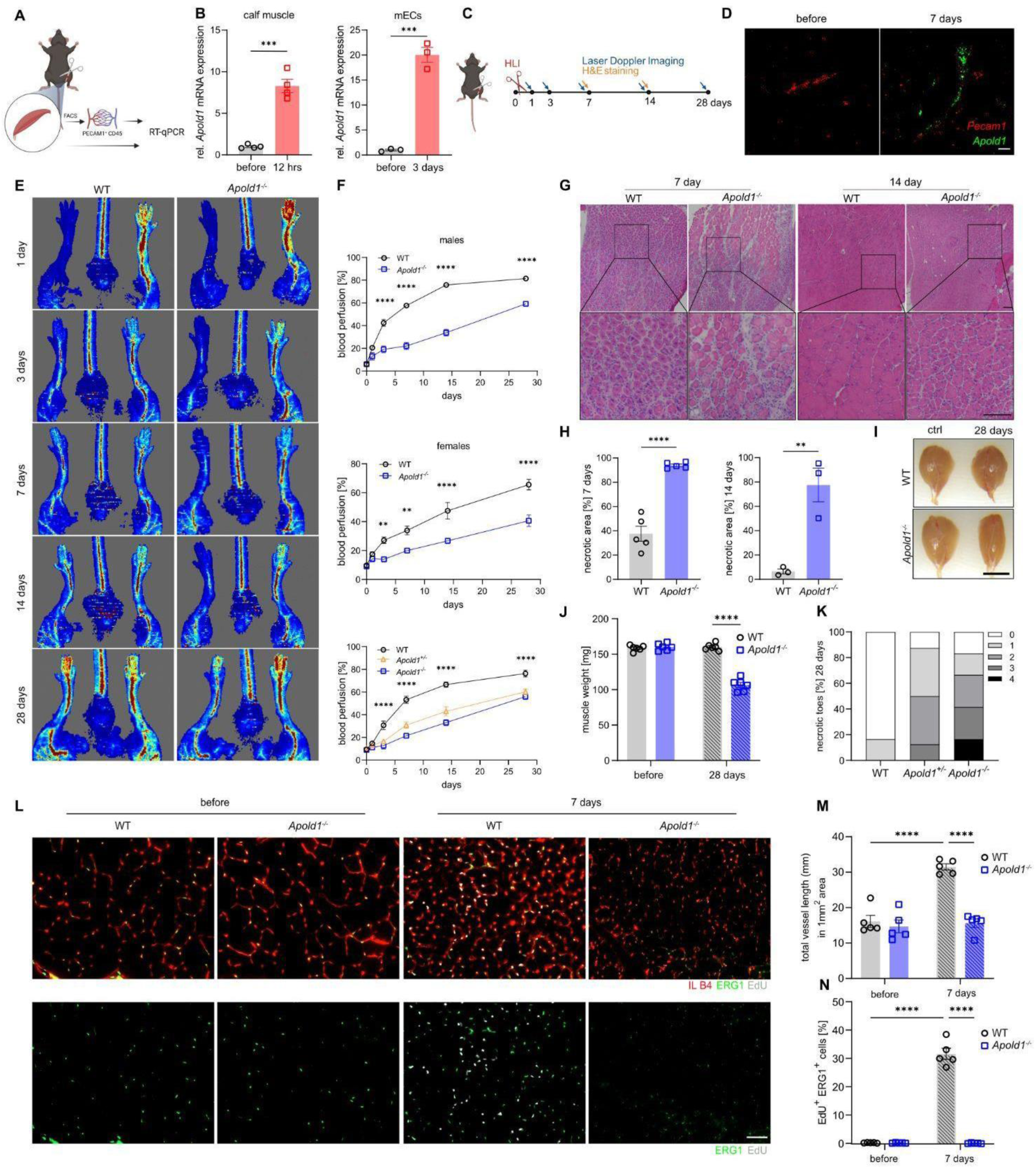
Impaired revascularization, angiogenesis, and EC proliferation after hindlimb ischemia in *Apold1^-/-^* mice. (A) Experimental design. (B) *Apold1* expression in qRT-PCR in whole calf muscle tissue 12 hours after hindlimb ischemia (left) and in muscle ECs (right) 3 days after hindlimb ischemia in WT mice (n=3-4 per group). (C) Experimental design. (D) Representative image of *in situ* hybridization for *Apold1* 7 days after ischemia in the ischemic muscle (ipsilateral) and the contralateral control muscle. Scale bar, 20 μm. (E) Representative images of blood perfusion measured by laser doppler imaging (LDI) 1, 3, 7, 14, 28 days after hindlimb ischemia in *Apold1^-/-^* and WT mice. (F) Time course quantification of blood perfusion across 28 days after hindlimb ischemia comparing recovery in *Apold1^-/-^* vs. WT males (n (WT/*Apold1^-/-^*) = 6/6), females (n (WT/Apold1^-/-^) = 6/6), and in a separate experiment in *Apold1^-/-^*, *Apold1^+/-^* and WT males (n (WT/*Apold1^+/-^*/*Apold1*^-/-^) = 6/6/4). (G) Representative hematoxylin-eosin (H&E) staining images of the triceps surae muscles at 7 and 14 days after hindlimb ischemia. Scale bar, 100 μm. (H) Quantification of necrotic area 7 and 14 days after hindlimb ischemia in WT and *Apold1*^-/-^ mice (n (WT/*Apold1^-/-^*) = 6/6). (I) Photographs of whole calf muscle isolated from control and ischemic leg of WT and *Apold1^- /-^* mice. Scale bar, 7 mm. (J) Whole calf muscle weight 28 days after hindlimb ischemia (n = 6 per group). (K) Measurement of number of necrotic toes 28 days after hindlimb ischemia (n = 8-12 per group). (L) Representative images of isolectin B4 (IL B4, red), Erg1 (green) and EdU (white) immunofluorescent images on gastrocnemius muscle cross sections of control and ischemic leg 7 days after hindlimb ischemia in WT and *Apold1^-/-^* mice. Scale bar, 50 μm. (M) Total vessel length in 1 mm^2^ area of muscle cross sections before and 7 days after hindlimb ischemia in WT and *Apold1^-/-^* mice (n=5 per group). (N) Percentage of EdU^+^ Erg1^+^ proliferating mECs at 7 days after induction of hindlimb ischemia (n=5 per group). Student’s t-test in B, H and J; Two-way ANOVA with Sidak’s multiple comparison test in F; Two-way ANOVA with Tukey’s multiple comparison test in M and N (**p<0.01; ***p<0.001; ****p<0.0001). The data shown are mean ± SEM.

Histological examination of H&E stained muscle sections revealed that *Apold1^-/-^* triceps surae muscles had much larger necrotic areas at 7 days post ischemia (Fig. 4G,H). Fourteen days after ischemia *Apold1^-/-^* mice still had large areas containing necrotic fibers, while WT muscles had almost completely recovered (Fig. 4G,H). In addition, while WT mice already showed clear signs of regeneration (evidenced by centrally nucleated fibers) at 7 days, *Apold1^-/-^* muscle only had low numbers of regenerating fibers, even at 14 days after ischemia (Fig. 4G). Further, impaired muscle recovery resulted in lower muscle weight in *Apold1^-/-^* mice 28 days after ischemia (Fig. 4I,J). We also noticed a higher frequency of necrotic toes in *Apold1^-/-^* mice and in haplo-deficient *Apold1^+/-^* mice (Fig. 4K). Finally, we measured vascular density in ischemic and (contralateral) control muscles on day 7 after hindlimb ischemia using isolectin B4 (IB4). While ischemic muscles in WT mice showed the typical increase in total vessel length, this response was completely abolished in *Apold1^-/-^* mice (Fig. 4L, M). Consistent with our observations in stroke, EdU labeling of proliferating mECs at 7 days following induction of ischemia revealed that *Apold1-*deficient mECs were unable to proliferate in response to ischemia (Fig. 4L,N).

### *Apold1* controls tumor angiogenesis

Because of the profound role of *Apold1* in regulating angiogenesis in pathological settings, we next asked whether *Apold1* controls tumor angiogenesis. We first used a published scRNAseq dataset and quantified *Apold1* expression in ECs of normal versus malignant lung tissue resected from 5 patients with untreated, non-metastatic lung tumors (Fig. 5A,B) [51]. Strikingly, we found that within ECs, *Apold1* expression was enriched in tumor ECs (tECs) compared to pulmonary ECs (pECs) (Fig. 5C). We confirmed these findings by re-analyzing a second scRNA-seq dataset, where freshly isolated human tECs and peritumoral non-tumor pECs (collected from the same individual) were analyzed from 1 large cell carcinoma, 4 squamous cell carcinomas, and 3 adenocarcinoma of treatment-naive patients [52]. Again, tECs expressed more *Apold1* than pECs (Fig. S3). To functionally test the role of *Apold1* in tumorigenesis, we implanted syngeneic B-16-F10 melanoma cells in WT and *Apold1^-/-^* mice and followed tumor growth over time (Fig. 5D). While WT mice showed exponential tumor growth, tumor growth was retarded in *Apold1^-/-^* mice leading to lower tumor volume and tumor weight at end-stage (Fig. 5E,F). Combined PECAM1 and ACTA2 staining showed similar vessel (PECAM1^+^) area with similar coverage by pericytes (ACTA2^+^) in the tumor tissue of both WT and *Apold1^-/-^* mice (Fig. 5G,I). However, closer investigation of the tumor vasculature showed that *Apold1^-/-^* mice had more vessels (Fig. 5J), but these vessels had a much smaller lumen (Fig. 5K), resulting in a dramatic reduction in lumen area per vessel in *Apold1^-/-^* mice (Fig. 5L). Functional assessment of vessel perfusion following intravenous injection with fluorescein-labeled Lycopersicon esculentum (tomato) lectin confirmed these data (Fig. 5 M-O). Finally, we injected EdU four hours before sacrificing the mice and subsequently FACS-sorted PECAM1^+^/CD45^-^ tumor ECs (Fig. 5P,Q). We found that loss of *Apold1* reduced the percentage of proliferating (Edu^+^) ECs leading to a lower fraction of ECs inside the tumor (Fig. 5R,S).

**Fig. 5.**
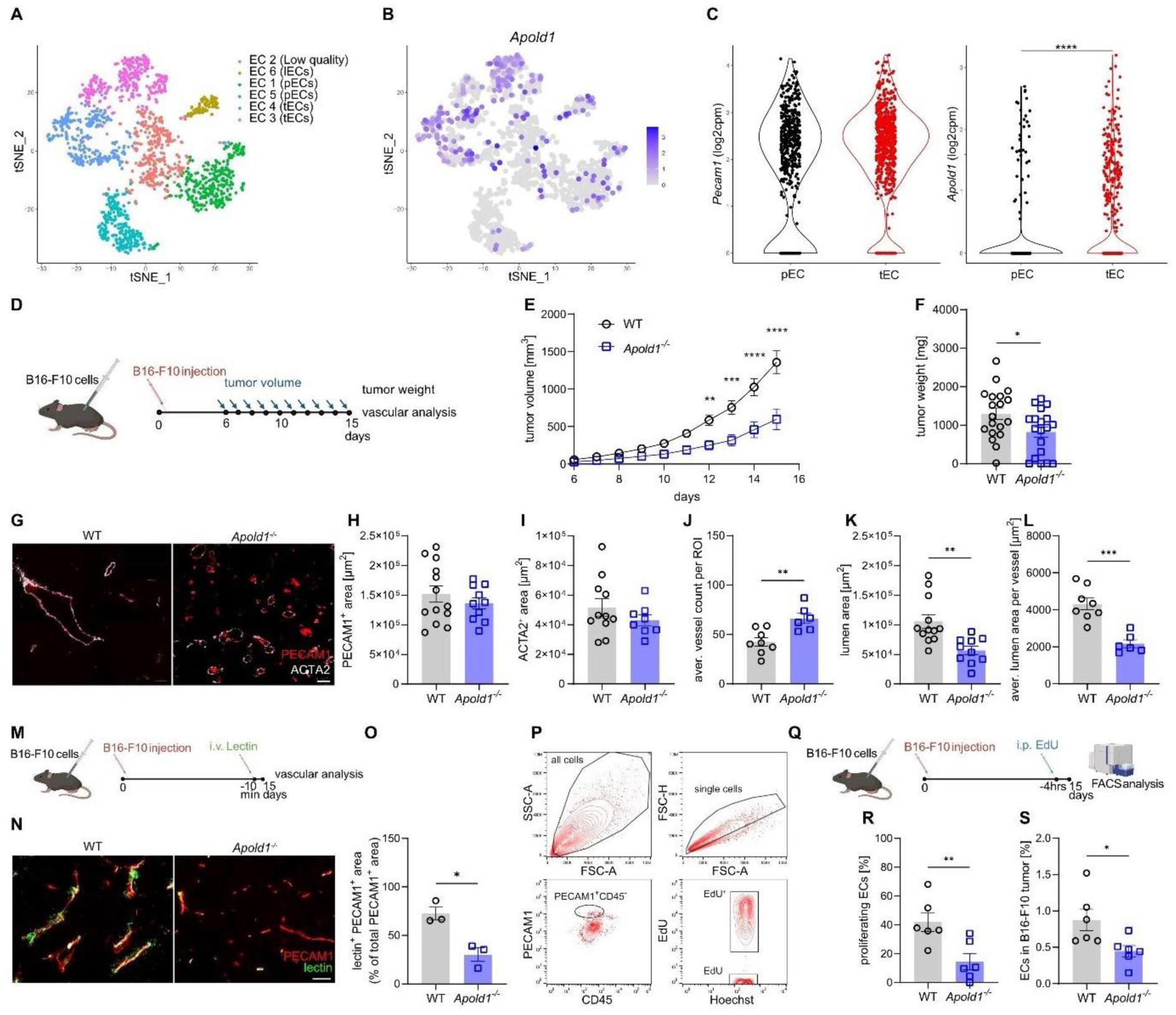
A*p*old1 is enriched in tumor ECs and loss of *Apold1* slows tumor growth. (A) t-Stochastic neighbor embedding (tSNE) plots of distribution of ECs in lung tissue resected from 5 patients with untreated, non-metastatic lung tumors reanalyzed [51]. (B) t-Stochastic neighbor embedding (tSNE) plots of distribution of *Apold1* in ECs in lung tissue reanalyzed [51]. (C) *Apold1* expression in tumor ECs (tECs) compared to pulmonary ECs (pECs) reanalyzed [51]. (D) Experimental design. (E) *In vivo* measurement of tumor volume in *Apold1^-/-^* and WT mice after injection of B16-F10 melanoma cells (n (WT/*Apold1^-/-^*) = 11/11). (F) Weight of isolated tumors 15 days after injection (n (WT/*Apold1^-/-^*) = 19/19). (G) Representative images of the vasculature in tumors isolated from WT and *Apold1*^-/-^ mice stained for PECAM1 (red) and ACTA2 (white). Scale bar, 100 µm. (H) Quantification of PECAM1^+^ area (n (WT/*Apold1^-/-^*) = 13/10) and (I) ACTA2^+^ area (n (WT/*Apold1^-/-^*) = 13/10). Scale bar, 100 µm. (J) Quantification of average vessel count per region of interest (ROI) (n (WT/*Apold1^-/-^*) = 8/6), (K) lumen size (n (WT/*Apold1^-/-^*) = 12/10), and (L) average lumen area per vessel (n (WT/*Apold1^-/-^*) = 8/6). (M) Experimental design. (N) Representative image of perfusion stained by injected fluorescein-labeled Lycopersicon esculentum (tomato) lectin (green) in PECAM1^+^ (red) in tumors isolated from WT and *Apold1*^-/-^ mice. (O) Percentage of lectin^+^PECAM1^+^ area of total PECAM1^+^ area (n (WT/*Apold1^-/^*) = 3/3). (P) Representative flow cytometric analysis of ECs (PECAM1^+^CD45^-^) and proliferating ECs (EdU^+^) in B16-F10 melanoma. (Q) Experimental design. (R) Percentage of proliferating ECs (EdU^+^) (n (WT/*Apold1^-/-^*) = 6/6) and (S) percentage of ECs (PECAM1^+^CD45^+^) in B16-F10 melanoma (n (WT/*Apold1^-/-^*) = 6/6). Student’s t-test in C, F, H, I, J, K, L, O, R, and S; Two-way ANOVA with Sidak’s multiple comparison test in E (*p<0.05; **p< 0.01; ***p< 0.001). The data shown are mean ± SEM.

Altogether, these results demonstrate a critical role for *Apold1* in angiogenesis and revascularization after hypoxic injury in the CNS and in the periphery, as well as during tumor growth. This is particularly striking when considering that disruption of many vascular genes involved in angiogenesis leads to severe developmental deficits, whereas *Apold1*-deficient mice are developmentally normal, suggesting that *Apold1* plays a role primarily during angiogenesis in pathological settings.

### Apold1 controls EC proliferation

To evaluate whether the ability of *Apold1* to control angiogenesis occurs via EC intrinsic mechanisms, we isolated and cultured primary ECs from *Apold1^-/-^* mice and wild-type littermates (Fig. 6A). We observed that ECs from *Apold1^-/-^* mice proliferated less in highly angiogenic cell culture medium (Fig. 6B,C). Moreover, spheroids created from ECs of *Apold1^-/-^* mice showed less sprouting than those from wild-type mice (Fig. 6D,E). To test whether *Apold1* is also required for cell proliferation in human umbilical vein endothelial cells (HUVECs), we knocked down *Apold1* using lentiviral transfection with short hairpin RNAs (shRNAs), which reduced *Apold1* expression by >50% compared to scrambled controls (Fig. 6F). Similar to our findings in *Apold1^-/-^* mECs, *Apold1* knockdown (Apold1-KD) in HUVECs reduced proliferation (Fig. 6G,H), and it also dramatically reduced sprouting length in a spheroid assay (Fig. 6I,J). This impairment was dependent on proliferation, since inhibiting proliferation using MitomycinC lowered sprouting similar to Apold1-KD and prevented any further decrease in sprouting after Apold1-KD (Fig. 6K). Indeed, scratch assays confirmed that migration into the scratch was not impaired after *Apold1*-KD (Fig. 6L,M).

**Figure 6.**
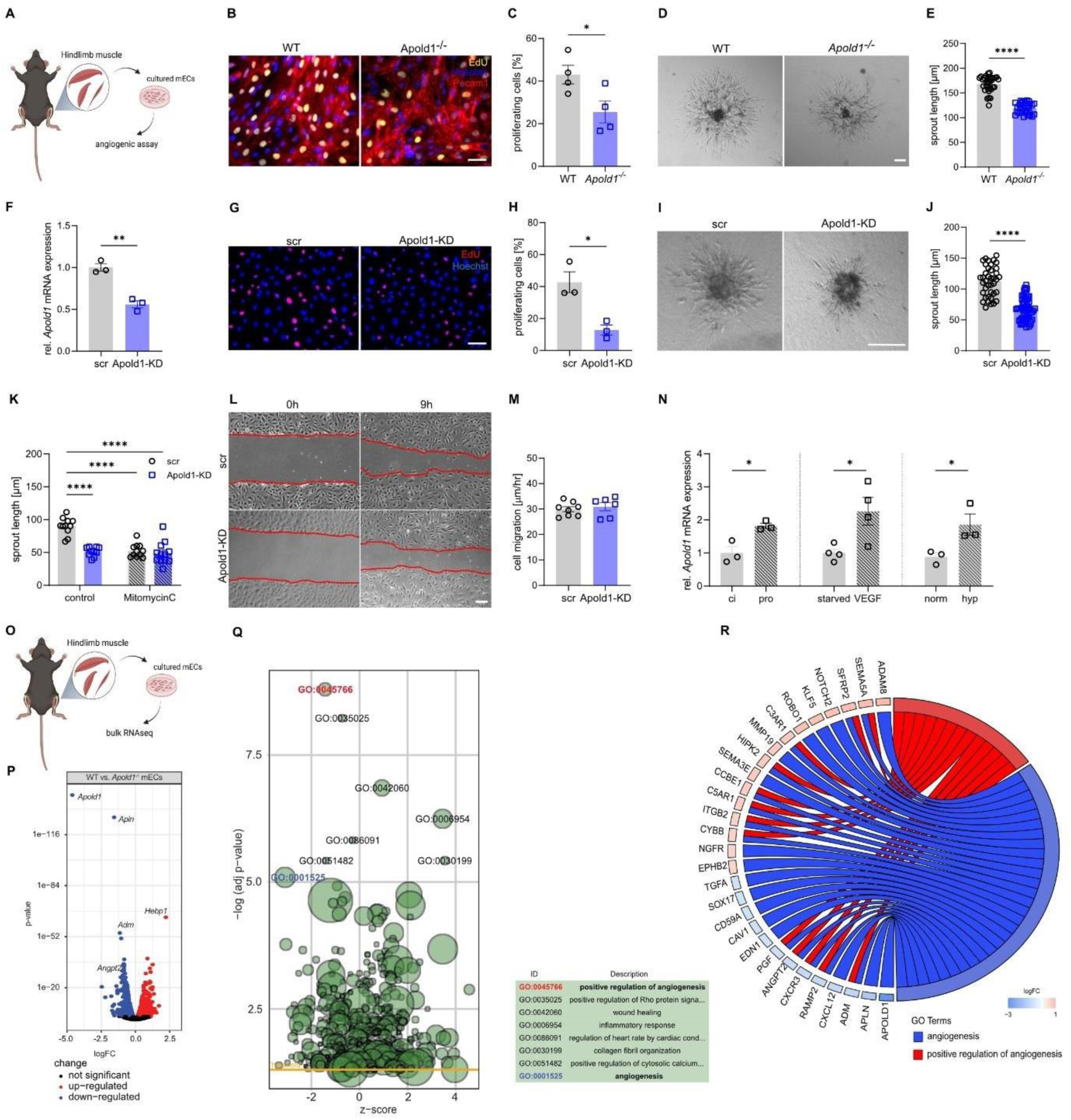
Angiogenesis is impaired in *Apold1*-deficient cells *in vitro*. (A) Experimental design. (B,C) Representative immunofluorescent images and quantification of three independent experiments of EdU (yellow) labeling of proliferating ECs isolated from skeletal muscle of WT and *Apold1^-/-^* mice, co-stained with Hoechst (blue) (n (WT/*Apold1^-/-^*)= 3/3. Scale bar, 100 μm). (D,E) Representative brightfield images and morphometric quantification of average sprout length of sprouting spheroids of mECs isolated from WT and *Apold1^-/-^* mice (n (WT/*Apold1^-/-^*) = 30/33. Scale bar, 100 μm). (F) *Apold1* knock-down efficiency in HUVECs treated with scrambled (scr) shRNAs or shRNAs against *Apold1* (Apold1-KD). (G, H) Representative immunofluorescent images and quantification of percentage of EdU (red) labeled proliferating HUVECS to all nuclei stained with Hoechst (blue) in scr and Apold1-KD HUVECs (n (WT/Apold1-KD) = 3/3. Scale bar, 100 μm). (I,J) Representative image and quantification of sprout length of spheroids of scr and Apold1-KD HUVECs (n (scr/Apold1-KD) = 30/33. Scale bar, 100 μm). (K) Quantification of sprout length in scr and Apold1-KD HUVECs treated in control conditions or MitomycinC (n (scr/Apold1-KD) = 10/10). (L,H) Representative images and quantification of cell migration in scratch assay (n (scr/Apold1-KD) = 7/6). (N) Relative *Apold1* expression in contact-inhibited (ci) and proliferative conditions (pro), after VEGF treatment and in hypoxic condition (0.1% O2) (n (scr/Apold1-KD) = 3/3). (O) Experimental design. (P) Volcano plots with red and blue dots showing significantly changed genes within 5% false discovery rate (FDR) determined by bulk RNAseq of cultured WT vs *Apold1^-/-^* mECs (n (WT/*Apold1^-/-^*) =3/3). (Q) Gene ontology analysis of bulk RNAseq of cultured and WT vs *Apold1^-/-^* mECs (n (WT/*Apold1^-/-^*) =3/3). (R) Overview of genes significantly differentially expressed within the GO terms angiogenesis (blue) and positive regulation of angiogenesis (red) in cultured WT and *Apold1^-/-^* mECs. Student’s test in C, E, F, H, J, M, and N; Two-way ANOVA with Tukey’s multiple comparison test in K (*p<0.05; **p< 0.01; ****p< 0.0001). The data shown are mean ± SEM.

Consistent with our *in vivo* data, we found that *Apold1* expression was higher under proliferating angiogenic conditions versus contact inhibition, when ECs are quiescent (Fig. 6N, left). Also, *Apold1* increased upon stimulation with VEGF (Fig. 6N, middle), while previous work already showed increased expression upon FGF2 [20]. Furthermore, since hypoxia is a known driver of angiogenesis, we confirmed that *Apold1* expression is increased in response to hypoxia in ECs [20, 30, 53, 54] (Fig. 6N, right). In fact, revisiting a published HIF1 ChIP-seq screen in HUVECs under normoxia or after 24hrs of hypoxia [55] revealed that hypoxia increases HIF binding to *Apold1* very similarly to the well-described HIF1 binding to VEGF (Fig. S4A). Additional analysis of this dataset showed that HIF1a knockdown abolished hypoxia-induced *Apold1* expression, further supporting *Apold1* regulation by HIF-1α (Fig. S4B). On the other hand, knocking down *Apold1* did not affect the regulation of HIF-responsive genes under hypoxia (Fig. S4C). Thus, *Apold1* can be activated by angiogenic growth factors as well as hypoxia. Altogether, *Apold1* is required to increase EC proliferation in response to a variety of angiogenic stimuli.

To subsequently explore how *Apold1* affects angiogenesis, we initially focused on autophagy, since it is a known regulator of endothelial function under pathological conditions [56] and because other apolipoproteins have been linked to autophagy before [57]. Moreover, autophagy was recently reported to be impaired upon *Apold1* knockdown in human dermal blood ECs, but we could not find any differences in LC3B nor p62 expression during normal culture conditions, nor under amino acid deprivation, a condition known to activate autophagy (Fig. S5A,B). We next performed RNA sequencing on primary ECs derived from *Apold1^-/-^* or WT ECs, which were briefly (16 hrs) cultured in angiogenic medium (Fig. 6O). We found 2255 genes to be differentially expressed between *Apold1^-/-^* and WT mECs (1041 up, 1246 down; FDR adj.p < 0.05; Fig. 6P; see list of differentially expressed genes in Supplementary Table2). These results are in striking contrast to the subtle gene expression differences observed under baseline conditions (Fig. 2I), and suggest that pro-angiogenic conditions unmask a profound functional impairment in *Apold1*-deficient ECs. In support of our findings, GO analysis showed that main top downregulated pathways were linked to angiogenesis and positive regulation of angiogenesis (Fig. 6Q). In fact, most genes included in those GO terms which were affected upon loss of *Apold1* all crucially control vascular development and many, when genetically removed, are associated with severe developmental phenotypes or even lethality. These data underscore the role of *Apold1* as a regulator of angiogenesis under pathological conditions.

## Discussion

In this study, we report that the vascular gene *Apold1* is dispensable for developmental angiogenesis but crucially contributes to ischemia-induced revascularization of the brain (stroke) and muscle (femoral artery ligation) as well as tumor angiogenesis by controlling EC proliferation. Despite the large phenotypes observed under such conditions, we did not observe a role for *Apold1* during developmental angiogenesis. *Apold1^-/-^ mice* were born at the expected Mendelian frequencies, showed normal development, and showed normal retinal angiogenesis. Furthermore, we did not find any differences in vascular density of adult brain nor muscle, and *Apold1^-/-^* mice behaved similar to their WT littermates. These data are remarkable since the constitutive deletion of angiogenic regulators often leads to severe developmental phenotypes or even lethality. In fact, there are only few genes such as *Placental Growth Factor* [58], *Vegfb* [59], *Ang2* [60] or *Robo4* [61] that upon constitutive deletion do not impair development but only affect angiogenesis in pathological conditions.

It is unclear why *Apold1* does not affect developmental angiogenesis while strongly restricting angiogenesis in several pathological settings. We showed that *Apold1* expression is activated by VEGF, which is known to activate an immediate early gene response [62], as well as by hypoxia. A previous RNAseq screen already identified *Apold1* to be part of a HIF-dependent angiogenic response downstream of YAP/TAZ in bone ECs [53]. We also found higher *Apold1* levels in muscle ECs during hindlimb ischemia, and in the brain upon stroke. In those conditions, deleting *Apold1* impaired angiogenesis. Previous reports however also observed higher *Apold1* expression during development [20], a condition not affected by *Apold1* deletion. What is interesting to note is that *Apold1* is also rapidly activated under specific conditions such as stress, which are not commonly associated with angiogenesis. This might indicate that the role of *Apold1* is not restricted to angiogenesis but contributes to the adaptive, homeostatic response to stress, consistent with its original description as an early immediate gene [20]. In such a scenario, *Apold1* might fine-tune the normal angiogenic response to quickly restore tissue homeostasis.

Given the high expression of *Apold1* during development, a previous report evaluated loss of *Apold1* in a mouse model of neonatal stroke. *Apold1-*deficient pups showed reduced angiogenesis after stroke and had impaired long-term functional recovery [26]. However, in adult mice, where *Apold1* expression is much lower, acute stroke caused similar size lesions and comparable functional impairment (24-72 hours after stroke) in *Apold1^-/-^* mice and wild-type controls [33]. In agreement with those observations, we found that stroke size was not different between genotypes, but, importantly, revascularization at the ischemic border was highly reduced in *Apold1^-/-^* mice. Because revascularization of the ischemic area is crucial for functional recovery in preclinical mouse models and patients after stroke [63–65], we speculate that loss of *Apold1* might impair functional recovery in stroke.

Although little is known about the possible role of *Apold1* in cancer, two studies have reported dysregulation of *Apold1* expression through DNA methylation. One study reported hypermethylation of *Apold1* in testicular germ cell tumors and in testicular embryonal carcinoma NT2 cells, lower *Apold1* expression in two types of testicular tumor (seminoma and embryonal carcinoma), and increased *Apold1* expression in response to treatment with the demethylating agent 5-azacytidine [66]. In contrast, another study found that *Apold1* DNA is hypomethylated and its expression strongly increased in two independent cohorts of patients with colorectal cancer [67]. Vascular heterogeneity, determined by tumor origin and type, is a hallmark feature of cancer and might explain these differences in *Apold1* regulation [14]. Therefore, we used single cell datasets to evaluate *Apold1* expression in human cancer. We found that *Apold1* expression is restricted to ECs and is higher in tumor ECs when compared to normal ECs [51, 52], confirming its relevance in humans. In a mouse model of melanoma (B16-F10), we found that subcutaneous tumor growth was reduced in *Apold1^-/-^* mice. Vessels were smaller, with very small lumens, leading to impaired tumor perfusion. Further experiments showed that ECs proliferated much less, leading to a lower fraction of ECs in the tumor. While these data require confirmation in other, more chronic tumor models, they suggest that inhibiting *Apold1* could prevent uncontrolled endothelial proliferation in pathological settings.

Interrogating several mouse as well as human single cell sequencing datasets, we consistently found that *Apold1* expression was restricted to vascular cells, predominantly ECs. Some datasets also revealed expression in pericytes and/or SMCs, but those cells were much less responsive to metabolic challenges, while endothelial *Apold1* was highly activated under such conditions. Since we used constitutive *Apold1^-/-^* mice in our experiments, we cannot rule out a contribution of perivascular cells to the observed phenotypes, nor can we exclude that *Apold1* controls angiogenesis by affecting the crosstalk between ECs and other cells. Nonetheless, our *ex vivo* and *in vitro* experiments all confirmed a strong cell-intrinsic role of *Apold1* inhibition leading to reduced endothelial cell proliferation, a key feature of angiogenic ECs. How *Apold1* controls proliferation remains an outstanding question. We found a concerted downregulation of many pro-angiogenic genes and growth factors in *Apold1^-/-^* conditions. It is not clear though whether this is directly linked to *Apold1* function, or secondary to the impaired proliferative capacity downstream of *Apold1*.

Previous work has localized APOLD1 to the membrane (presumably focal adhesion sites) [20, 31] or at Weibel Palade bodies [31]. Unfortunately, we did not find any commercially available antibody that specifically labeled APOLD1 in WT versus *Apold1^-/-^* ECs, nor upon the use of overexpression/knock-down approaches in human ECs using Western blot as well as immunohistochemistry (data not shown). In addition, we did not observe changes in autophagy. Likely, the role of APOLD1 is contextual, and is dependent on the endothelial source, the growth state of the cell, the stimulus that induces APOLD1 function, and the specific culture conditions. For instance, Stritt et al. used confluent monolayers to evaluate the role of APOLD1 [31], while our experiments specifically aimed to study APOLD1 under angiogenic (non-confluent), growth factor stimulated conditions.

Taken together, we here show that *Apold1* is dispensable for developmental angiogenesis, but that it controls ischemia-induced revascularization of the brain (stroke) and muscle (femoral artery ligation), and regulates pathological tumor angiogenesis, probably largely by controlling EC proliferation. The absence of a developmental phenotype, the observation that *Apold1* is activated under ischemic and pro-angiogenic conditions, and the fact that its expression is restricted to vascular endothelial cells, could make it an interesting target for future therapeutic interventions to either enhance vascular repair or to restrict tumor vascularization.

## Data availability

The sequencing data generated in this study have been deposited in the Gene Expression Omnibus database under accession codes GSE217694.

## Authorship contributions statement

ZF, RA, JB, and KDB contributed to the study conception and design. ZF and RA performed and analyzed most experiments. AAB, LVZ, AEM, PLG performed bioinformatic analysis of RNAseq datasets. RR, RWe, RWa performed and analyzed stroke experiments. RWa helped with RNAseq. HD performed and analyzed retina experiments. JZ, TD, assisted with hindlimb ischemia surgeries and analysis. OS performed behavior experiments. ST, MES, JB, and KDB supervised the experiments and provided funding. The manuscript was written by ZF, RA, JB, and KDB. All authors read and approved the final manuscript.

## Acknowledgements

KDB is endowed by the Schulthess foundation. The lab of KDB is funded by the ETH Zurich and SNSF Grant 31003A_208041. The lab of JB is funded by the ETH Zurich, the ETH Project Grant ETH-20 19-1, SNSF Grant 310030_172889, Swiss 3R Competence Center, Botnar Research Centre for Child Health, Multi-Investigator Project. Apold1^-/-^ mice were generously provided by Prof. Paul Worley. We thank Han-Yu Lin for skilled technical assistance, and Prof. Sonia Tugues Solsona for thoughtful discussions. We thank the ETH Zurich Flow Cytometry Core Facility for the assistance.

## Methods

### RNA sequencing and data analysis

#### Sequencing

For the RNA sequencing of samples from FACS sorted mECs incubated under angiogenic conditions for 16h the following protocol was used: The TruSeq stranded RNA kit (Illumina Inc.) was used according to the manufacturer’s protocol. The mRNA was purified by polyA selection, chemically fragmented and transcribed into cDNA before adapter ligation. Single-end (100nt) sequencing was performed with Illumina Novaseq 6000. Samples were all run on the same lane and demultiplexed. A sequencing depth of ∼20M reads per sample was used.

For RNA sequencing of FACS sorted mECs without prior incubation, cDNA conversion was performed using the Smart-seq2 protocol [68] using 800 pg mRNA as input. Library preparation and paired-ended sequencing (150 bp) was performed by Novogene (Cambridge, UK) on an Illumina Novaseq 6000 machine. A sequencing depth of ∼20M read pairs per sample was achieved.

#### Analysis

Adapters were trimmed using cutadapt [69] (v 1.15) with a maximum error rate of 0.05 and a minimum length of 15. Kallisto [70] (v0.44.0) was used for pseudo alignment of reads on the transcriptome level using the genecode.vM17 assembly with 30 bootstrap samples. For single-end samples an estimated fragment length of 200 ± 20 was used. For differential gene expression (DGE) analysis we aggregated reads of protein coding transcripts on the gene level and used R (v. 4.0.3) with the package “edgeR” (v 3.32.1) [71] for analysis. A filter was used to remove genes with low expression prior to DGE analysis. EdgeR was then used to calculate the normalization factors (TMM method) and estimate the dispersion (by weighted likelihood empirical Bayes). For two group comparisons the genewise exact test was used. For multiple testing correction the Benjamini-Hochberg false discovery rate (FDR) method was used. GO analyses on significant genes vs. the background of all tested genes were performed using the R package “topGO” (v 2.42.0) with a node size of 10 and a fisher p-value cutoff of 0.05 as determined by the algorithm “weight01”. The *mus musculus* mapping "org.Mm.eg" was used across all three categories (biological processes, molecular function and cellular component).

#### Analyses of Apold1 expression and HIF binding based on Mimura et al (2012) [55]

Microarray data produced following both a hypoxic time series and siRNA knockdowns were obtained from the NCBI gene expression omnibus (accession: GSE35932) published by Mimura et al., 2012 [55]. To determine the relative expression values, “RMA” was used for background correction, “qspline” for normalisation and “liwong” as a summary method. Probes corresponding to APOLD1 and VEGFA were identified using the hgu133plus2.db R package (version: version 3.2.3). Four probes were identified for VEGFA from which the mean values were calculated to represent the relative expression level of the gene.

Raw Chip-seq data for the identification of HIF1 binding sites in HUVECS for both normoxic and hypoxic conditions were obtained via the accession GSE39089 [55]. Reads were trimmed using trimmomatic [72] and aligned to the GRCh38 genome using bowtie2 [73] with default parameters. Coverage tracks were normalized using Counts Per Million (CPM) reads mapped, and visualization was made with the *epiwraps* package.

#### Analyses of single-cell RNA-seq data from muscle tissue [36]

Single cell RNA-seq data of Mononuclear cells from hindlimb skeletal muscle in wild type mice was obtained from the GEO database via the accession GSE110878 [36]. All single cell analyses and visualization was carried out using Seurat (version 3.1.0). The two data sets available for uninjured wild-type mice were integrated using Seurat’s standard workflow. Integration anchors were identified following canonical correlation analysis for dimension reduction. T-SNE was applied to the integrated dataset for visualization and cluster identification. The makers used to assign cell-type identities to each cluster are shown in Table 3. These labels were chosen as to reflect those findings in the original Giordani paper and to distinguish APOLD1 expressing cell types [36].

#### Analyses of brain single-cell RNA-seq datasets

For the dataset from Wu et al., 2017 [39] (Fig 1J-K), the re-analysis described in Floriou-Servou et al., 2021 [74] was used. For the dataset from von Ziegler et al., 2022 [41] (Fig 1L-M), the processed data was obtained from the github repository of the original publication, and vascular cells were reassigned to more specific categories in a supervised fashion. The counts of markers for SMCs, ECs and pericytes (see Table 3) were summed for each cell type, and cells which did not have at least twice as much signal for markers of one cell type as for the others were excluded as ambiguous cells. Cells were then assigned to the cell type with the maximum signal.

For the dataset from Zeisel et al, 2018 [38] (Fig. S2), we downloaded the L5 dataset from http://mousebrain.org/adolescent/downloads.html, and used the TaxonomyRank4. To perfect the vascular cell types, we isolated cells that had been assigned to cell types containing any of the words smooth muscle, pericytes, endothelial or vascular, and re-analyzed them separately using *scran* 1.18.7 [75]. We used 20 components of a PCA based on the 2000 most expressed genes, followed by Louvain clustering on the KNN graph. Clusters were then manually linked to cell types using the markers from Table 3. For other cell types, the original annotation was used.

### Postnatal retinal angiogenesis model

To assess postnatal retinal angiogenesis, WT and *Apold1^-/-^* pups were sacrificed at P6. Thereafter, eyes were harvested for retinal dissection and analysis of blood vessel outgrowth. Different parameters were automatically quantified with the angiogenesis plug in tool of Image J (NIH).

### Stroke induction

Mice were anesthetized using isoflurane (5% induction, 1.5-2% maintenance; Attane, Provet AG). Analgesic (Novalgin, Sanofi) was administered 1 d prior to the start of the procedure via drinking water. A photothrombotic stroke to unilaterally lesion the sensorimotor cortex was induced on the right hemisphere, as previously described [48, 76]. Briefly, animals were placed in a stereotactic frame (David Kopf Instruments), the surgical area was sanitized, and the skull was exposed through a midline skin incision. A cold light source was positioned over the right forebrain cortex (anterior/posterior: −1.5–/+1.5 mm and medial/lateral 0 –/+2 mm relative to Bregma). 5 min prior to illumination, Rose Bengal (10 mg/ml, in 0.9% NaCl, Sigma) was injected intact skull for 8 min (mild stroke) or 10.5 min (severe stroke). To restrict the illuminated area, an opaque template with an opening of 3 x 1.5 mm (mild stroke) or 3 × 4 mm (severe stroke) was placed directly on the skull. The wound was closed using a 6/0 silk suture and animals were allowed to recover. For postoperative care, all animals received analgesics (Novalgin, Sanofi) for at least 3 days after surgery.

### Stroke volume quantification

Stroke volume was calculated from coronal brain sections stained with NeuroTrace 640/660 (ThermoFischer). Brain sections at six defined landmarks (2.5, 1.5, 0.5, −0.5, −1.5, −2.5 mm in relation to bregma) were analyzed for depth of the cortical lesion. The dorso-ventral, medio-lateral, and anterior-posterior stroke extent was then used to modulate a precise ellipsoid with the coordinates relative to Bregma.

### RNA extraction

RNA extraction of brain tissue was carried out using Quiagen RNeasy kit according to the manufacturer’s recommendations. qPCR was performed using SYBR green kit (iTaq Universal SYBR Green Supermix from Biorad) containing 0.5 µM of each primer with the following cycling conditions (hold stage: 95 °C, 10 min, 1 cycle; PCR stage (95°C, 15 s, 60 °C 1 min; 95 °C 15 s, 40 cycles; Melting curve (95 °C, 15 s, 60 °C, 1 min).

### EdU analysis

To label proliferating vascular endothelial cells mice received three consecutive i.p. injections of 5-ethynyl-2’-deoxyuridine (EdU, 50 mg/kg body weight, ThermoFischer) on day 6, 7 and 8 after stroke. EdU incorporation was detected 21 days after stroke using the Click-it EdU Alexa Fluor 647 Imaging Kit (ThermoFischer) on 40 µm free floating coronal sections. We quantified the number of EDU^+^ PECAM1^+^ cells per mm2 stroked brain tissue.

### Vascular analysis

Vascular quantification including vessel area fraction, length, branching was performed based on an automated script previously established [77, 78]. Briefly, for area fraction, the area covered by PECAM1 was quantified using ImageJ after applying a constant threshold. The vascular length was quantified using the “skeleton length” tool and number of branches was assessed using the “analyze skeleton” tool. For the vascular length and branches, results were normalized per mm^2^ of brain tissue.

### Behavioral characterization

#### Open-field testing

Open-field testing took place inside sound insulated, ventilated multiconditioning chambers (TSE Systems Ltd, Germany). The open-field arena (45 cm (l) 45 cm (w) 40 cm (h)) consisted of four transparent Plexiglas walls and a light gray PVC floor. Animals were tested under yellow light (4 Lux across the floor of the open field) with 60–65 dB of white noise playing through the speakers of each box. An infrared light also illuminated the boxes so that an infrared camera could be used to record the tests. Prior to testing each animal, the entire open-field arena was cleaned using 10 ml/l detergent (For, Dr. Schnell AG). The room housing the multiconditioning chambers was illuminated with red LED lights (637 nm). Animals were removed from their home cage by the tail and placed directly into the center of the open field. The doors of the conditioning chamber were then swiftly closed. Tracking/recording was initiated upon first locomotion grid beam break. All open-field tests were 10 minutes in duration.

### Light-Dark Box

Light-Dark box testing took place inside sound insulated, ventilated multiconditioning chambers (TSE Systems Ltd, Germany). The Light-Dark box (internal dimensions: 42.5 cm (l) 29.5 cm (w) 24.5 cm (h) (dark compartment 15cm (l) 29.5cm (w) with a centered square opening 6cm x 6cm) consisted of both transparent and infra-red permeable black Plexiglas walls and a light gray PVC floor. Animals were tested under white light (200 Lux across the floor of the light compartment) with 60–65 dB of white noise playing through the speakers of each box. An infrared light also illuminated the boxes so that an infrared camera could be used to record the tests. Prior to testing each animal, the entire arena was cleaned using 10 ml/l detergent (For, Dr. Schnell AG). The room housing the multiconditioning chambers was illuminated with red LED lights (637 nm). Animals were removed from their home cage by the tail and placed directly into the center of the light compartment. The doors of the conditioning chamber were then swiftly closed. Tracking/recording was initiated upon first locomotion grid beam break. All Light-Dark box tests were 10 minutes in duration.

### Hindlimb ischemia model

Hind-limb ischemia experiments were performed as described before [79]. Briefly, mice were anesthetized with isoflurane, the hind limb was shaved, and, following a small incision in the skin, both the proximal end of the femoral artery and the distal portion of the saphenous artery were ligated. The artery and all side-branches were dissected free; after this, the femoral artery and attached side-branches were excised. Immediately after surgery, perfusion was measured by Laser Doppler Imaging of plantar regions of interest (Moor Instruments Ltd, Axminster, Devon, England) and calculated as ratio of left (ligated) versus right (unligated) values. To label proliferating cells, an intraperitoneal injection of 5-ethynyl-2’-deoxyuridine (EdU) (E10187, Thermo Fischer Scientific) solution (5 mg/ml in saline, 10μg per gram body weight) was performed 7 hours before sacrificing the mice.

### Isolation of primary muscle endothelial cells (mECs)

<u>mEC isolation for mRNA analysis</u> was performed as described before [37]: Mice were euthanized and calf muscles from different groups were immediately dissected, and superficial big vessels were carefully removed. Muscles were minced in a petri dish on ice with a scalpel until a homogeneous paste-like mash was formed. Thereafter, the mashed muscle was enzymatically digested in digestion buffer containing 2 mg/ml Dispase II (D4693, Sigma-Aldrich, Steinheim, Germany), 2 mg/ml Collagenase IV (17104019, Thermo Fisher Scientific, Zurich, Switzerland), 2 mM CaCl2 and 2% BSA in PBS at 37°C for 10 min, with gentle shaking every 3 min. The reaction was stopped by immediately adding an equal volume of ice cold HBSS containing 20% FBS and the suspension was passed through a 70-μm cell strainer (#431751, Corning, New York, USA) then 40-μm cell strainer (#431750, Corning, New York, USA) to remove tissue debris. Cell suspension was centrifuged at 500 g for 5 min at 4 °C, then the pellet was washed with ice cold HBSS (+20% FBS) followed by a centrifugation at 400 g for 5 min in 4 °C. Next, the cell pellet was re-suspended in antibody medium (EGM2 CC-3162, Lonza, Basel, Switzerland) with anti-mouse CD31 (PECAM1) PE antibody (1:400) (553373, BD Biosciences, Basel, Switzerland) and anti-mouse CD45 PerCP antibody (1:400) (557235 BD Biosciences, Basel, Switzerland) and placed on ice for 20 min in the dark. Before sorting, the cell suspension was washed in FACS buffer (1xPBS+1%BSA) and centrifuged at 400 g for 5 min, 4 °C, then the washed cell pellet was re-suspended in FACS buffer containing cell viability dye, SYTOX^TM^ blue (1:1000) (S34857, Thermo Fischer Scientific, Zurich, Switzerland). Viable endothelial cells (PECAM1^+^, CD45^-^, SYTOX^TM^ blue^-^) were sorted by a FACS Aria III (BD Bioscience) sorter. 200,000 events (ECs) were directly sorted (70 μm nozzle) into 700 μl RNA lysis buffer, and RNA extraction was performed by RNeasy Plus Micro Kit (74034 QIAGEN).

#### mEC isolation for culturing

Whole skeletal muscle tissues from hind-limb were dissected and digested as described above. After a series of centrifugation and washing steps, the heterogeneous cell population was resuspended in EC culture medium and seeded in collagen type I (125-50, Sigma)-coated plates. Due to the higher expression of P-glycoprotein in ECs compared to other skeletal muscle cells, mECs were selected by adding 4 μg/ml puromycin (P8833, Sigma-Aldrich, St. Louis, USA) to the medium overnight. After 7 days in culture, the purity of mECs was determined by PECAM1 fluorescence staining and only cultures containing at least 85% of the cells positive for PECAM1 were used for further experiments.

### In-situ Hybridization

For in-situ hybridization, the RNAscope® Multiplex Fluorescent Assay v2 Kit (Cat.# 323110; Advanced cell diagnostics, Newark, United States) and probes (Tab. 3) were employed. The instructions were followed in accordance to the manufacturer’s manual. The procedure briefly described: <u>Tissue preparation for RNAscope</u>. Tissue were dissected and left in 4 % PFA until the tissue sunk to the bottom. Same procedure was repeated in 10 %, 20 % and 30 % sucrose (Cat.# 573113; MilliporeSigma, Burlington, United States). Samples were frozen in the fume of liquid nitrogen and stored at -80°C. Before sectioning (14 µm on Superfrost microscope slides), samples acclimatized in the cryostat (Leica CM1950 cryostat) at -20°C for 60 min and were embedded in OCT.

#### RNAscope pre-treatment

The sections were first washed in 1 x PBS for 5 min to remove OCT and treated with RNAscope Hydrogen Peroxide for 10 min at RT and washed in distilled water twice. For target retrieval, the slides were placed into slightly boiling (90-95°C) RNAscope 1 x Target Retrieval Reagents for 5 min. Slides were washed in distilled water and 100 % ethanol and air-dried at RT. After marking the slides with a hydrophobic pen RNAscope protease III was applied to the sections and incubated in the HybEZ Oven at 40°C for 30 min. Slides were washed in distilled water.

#### RNAscope

The hybridization step occurred by mixing pre-warmed probes (Tab.1; water bath at 40°C) as followed and adding the mix to the cover slides and incubated at 40 °C for two hours: Slides were washed twice with 1 x wash buffer for 2 min. Excess liquid was removed and the RNAscope Multiplex FL v2 Amp1 was added to the sections and incubated for 30 min. After washing the slides twice for 2 min with 1 x washing buffer, RNAscope Multiplex FL v2 Amp2 was added and again incubated at 40°C for 30 min. Same washing step occurred and RNAscope Multiplex FL v2 Amp 3 was applied to the slides and incubated at 40°C for 15 min. The slides were washed twice in 1 x wash buffer for 2 min and excess liquid was removed. RNAscope Multiplex Fl v2 Hrp-C1 was added to slides and incubated for 15 min at 40°C. Meanwhile, each opal dye in DMSO (520 nm, 570 nm, 650 nm; Cat#. NEL810001KT; PerkinElmers, Waltham, United States) was mixed 1:2000 in TSA buffer. After another washing step, opal 520 was applied to the sections and incubated for 30 min at 40°C. Once more, the slides were washed in 1 x washing buffer and blocked with RNAscope Multiplex FL v2 HRP blocker for 15 min at 40°C. Slides were washed twice in 1 x washing buffer. This procedure was repeated for RNAscope-Multiplex FL v2 HRP-C2 and Opal 570, followed by RNAscope-Multiplex FL v3 HRP-C3 and Opal 650. Lastly, sections were incubated with DAPI for 30 sec, mounted with DAKO and dried overnight in the dark at RT. Slides were stored at 4 °C in the dark.

### Cell culture

Isolated primary mouse skeletal muscle endothelial cells (mECs) and commercially purchased human umbilical vein endothelial cells (HUVECs) from pooled donors (C-12203, PromoCell, Heidelberg, Germany) were routinely cultured in a 1:1 ratio of M199 ((11150059, Thermo Fisher Scientific) supplemented with 20% fetal bovine serum (FBS) (10270-106, Thermo Fisher Scientific), 30 mg/L endothelial cell growth factor supplements (EGCS) (E2759, Sigma-Aldrich), 10 U/ml heparin (H3149 Sigma-Aldrich) and 1% Penicillin-Streptomycin (10,000 U/ml) (15140122, Thermo Fisher Scientific) and Endopan 3 (P04-0010K, PAN BIOTECH, Aidenbach, Germany) (denoted as M+E). Murine B-16 (F10) cells were purchased from ATCC® (CRL-6475™, Molsheim, France) and cultured in Dulbecco’s Modified Eagle Medium (DMEM) (Thermo Fisher, 41965039, Zurich, Switzerland) containing 10 % fetal bovine serum (FBS) and 1 % pen strep (PS) (100 IU/ml penicillin and 100 μg/ml streptomycin). Primary mECs were only used until passage (P)1, HUVECs were used between P1 and P5. Cells were routinely maintained in 5% CO2 and 95% air at 37 °C, and regularly tested for mycoplasma.

### RNA extraction and quantitative RT-PCR

RNA of cultured mECs and HUVECs was extracted using PureLink™ RNA Mini Kit (12183020, Thermo Fischer Scientific). For RNA isolation from muscle tissues, 100 mg muscles were quickly dissected and homogenized in 1 ml Trizol, after 5 min incubation, 200 µl of chloroform was added and spined down at 1,200 x g for 15 min at 4 °C. Then transfer the transparent upper phase to a new tube and add equal volume of 70 % ethanol. Transfer to a RNeasy Mini spin column and process as described above. RNA purity and concentration were assessed via a spectrophotometer (Spark 10M, Tecan). RNA was reverse-transcribed to cDNA by High Capacity cDNA Reverse Transcription Kit (Thermo Fisher Scientific, 43-688-13). A SYBR Green-based master mix (Thermo Fisher Scientific, A25778) was used for real-time qPCR analysis with primers listed in Table 2. To compensate for variations in RNA input and efficiency of reverse-transcription, 18S was used as a housekeeping gene. The delta-delta CT method was used to normalize the data.

**Table 1.** Overview of RNAscope probes. Listed are all employed probes purchased from Advanced cell diagnostics and their according gene, the species, the relative volume used and their Cat.#.

| Channel | Gene | Species | Relative volume | Cat.# |
| --- | --- | --- | --- | --- |
| 1 | Apold1 | Mm | 50 x | 418161 |
| 2 | Pecam1 | Mm | 1 x | 316721-C2 |
| 3 | Acta2 | Mm | 1 x | 319531-C3 |

**Table 2.**
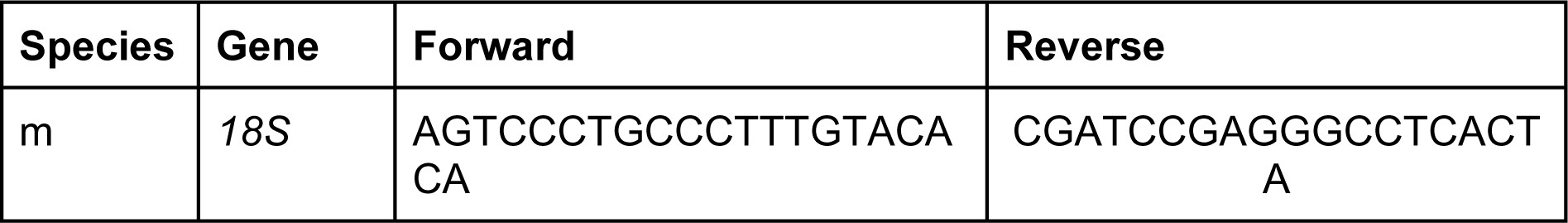
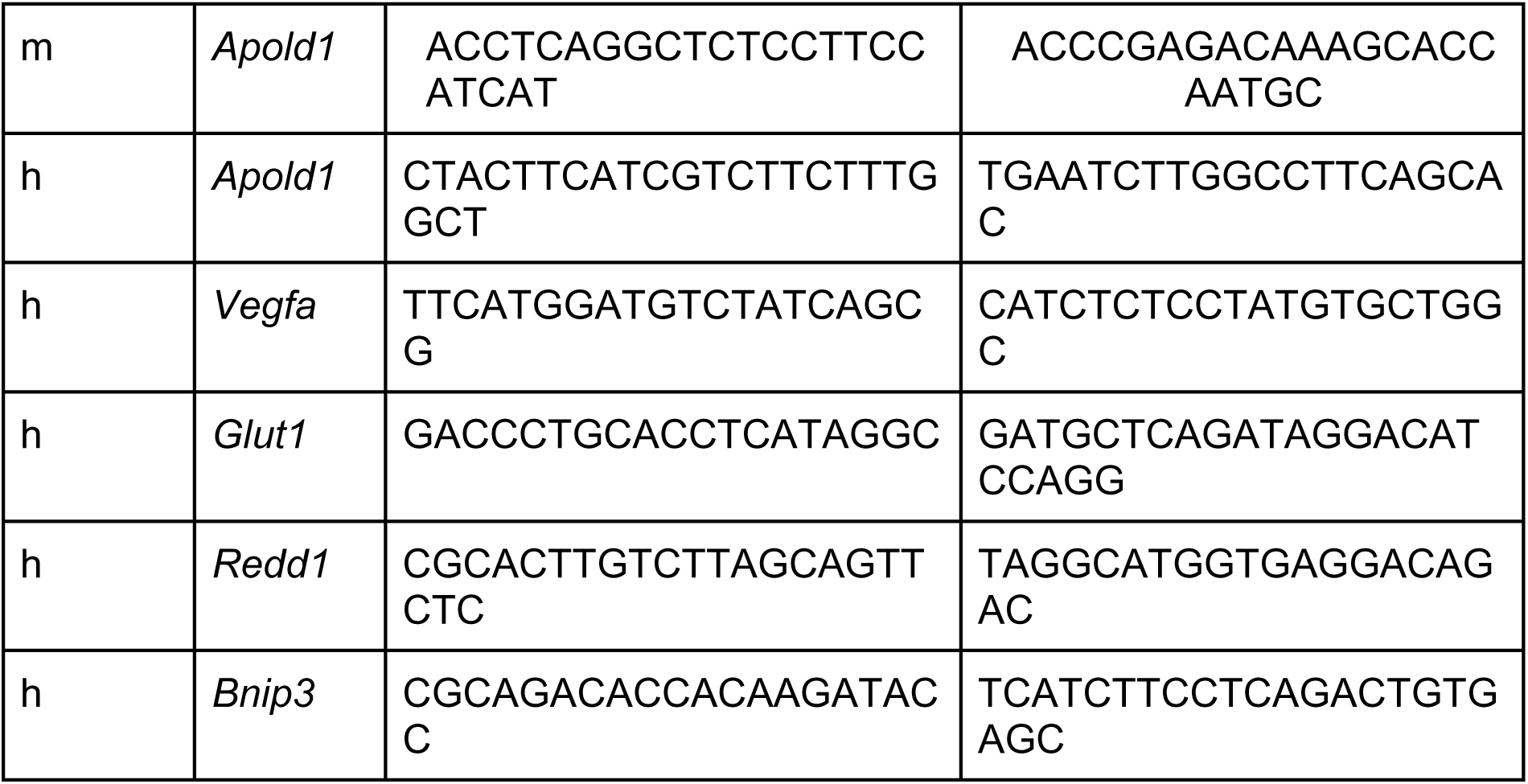
Forward and reverse primer sequences for murine (m) and human (h) primers used for RT-qPCR.

**Table 3:**
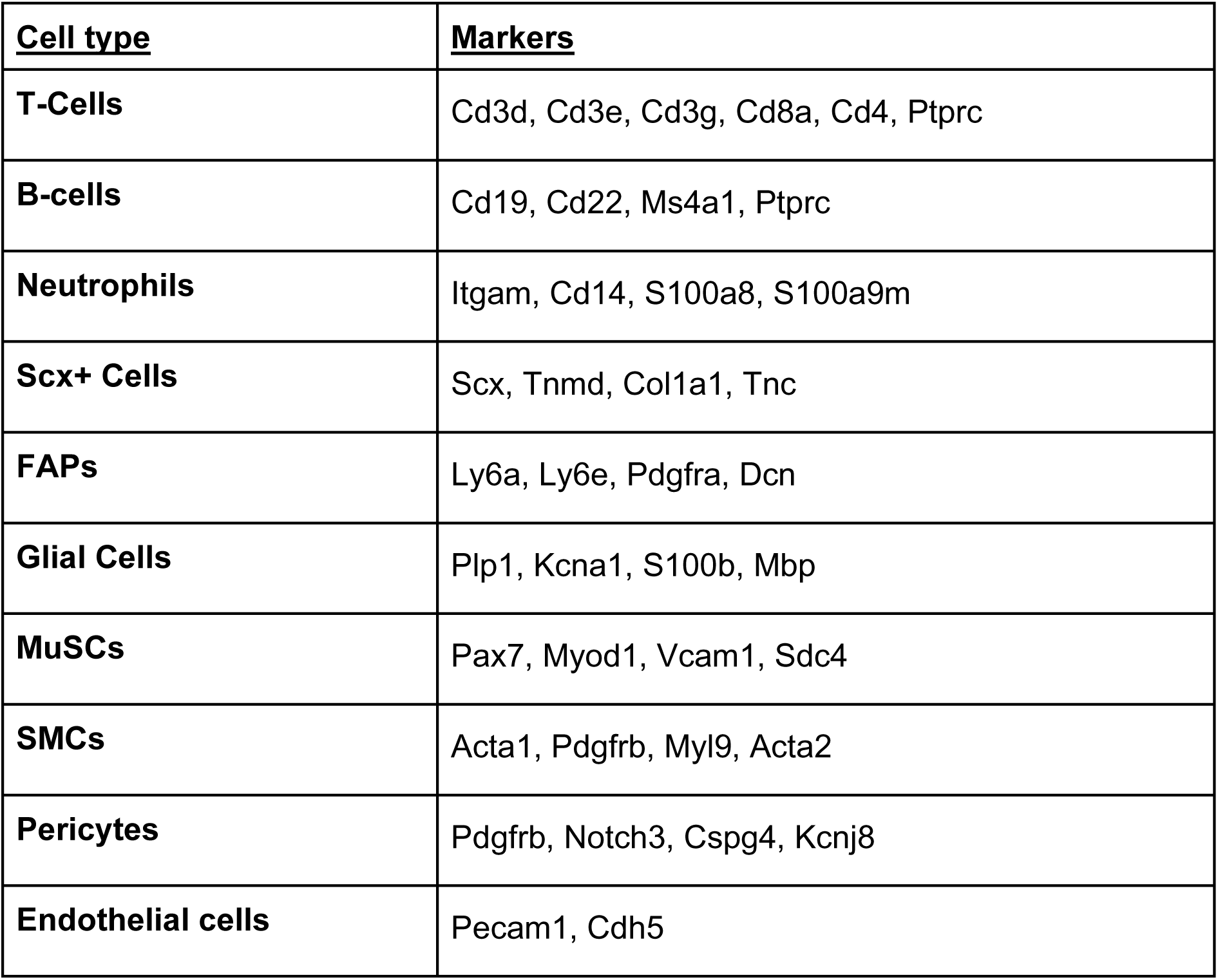
Overview of cell types and their respective markers used for cell annotation.

| **<u>Cell type</u>** | **<u>Markers</u>** |
| --- | --- |
| **T-Cells** | Cd3d, Cd3e, Cd3g, Cd8a, Cd4, Ptprc |
| **B-cells** | Cd19, Cd22, Ms4a1, Ptprc |
| **Neutrophils** | Itgam, Cd14, S100a8, S100a9m |
| **Scx+ Cells** | Scx, Tnmd, Col1a1, Tnc |
| **FAPs** | Ly6a, Ly6e, Pdgfra, Dcn |
| **Glial Cells** | Plp1, Kcna1, S100b, Mbp |
| **MuSCs** | Pax7, Myod1, Vcam1, Sdc4 |
| **SMCs** | Acta1, Pdgfrb, Myl9, Acta2 |
| **Pericytes** | Pdgfrb, Notch3, Cspg4, Kcnj8 |
| **Endothelial cells** | Pecam1, Cdh5 |

### Subcutaneous tumor model

Mice at the age of 7-10 weeks were subcutaneously injected cultured murine B-16 (F10) melanoma cells (1 million cells per mouse) proximate under the left forelimb. The mice were monitored over the next 15 days to detect and quantify tumor growth. Tumor diameters were measured by caliper and the size was calculated using the formula: Tumor volume (mm^3^) = d^2^x D/2 where d and D are the shortest and longest diameter in mm, respectively. After two weeks, mice were anesthetized and sacrificed by transcardial perfusion with 2 % paraformaldehyde (PFA, Cat.# J61984 Alfa Aesar, Haverhill, United States). Tumors were dissected and fixed with 4 % PFA for 24-48 hours and frozen in the fume of liquid nitrogen. Samples were stored at -80°C.

<u>Tumor</u> <u>vessel perfusion</u> was quantified on tumor cryosections following intravenous injection of 0.05 mg fluorescein-labeled Lycopersicon esculentum (Tomato) lectin (FL-1171, Vector laboratories, Brussels, Belgium) in tumor-bearing mice (tumors were harvested 10 min after injection). The perfused area was defined as the lectin^+^ PECAM1^+^ area expressed as a percentage of the total PECAM1^+^ area (NIH Image J software). Tumor perfusable area was analyzed by measuring the total vessel lumen area (i.e., sum of the lumen area of all vessels), and expressing it as a percentage of the total tumor area.

### Knock down and lentiviral particle production

Multiple GIPZ Lentiviral shRNAs target human *APOLD1* (V2LHS_117002; V2LHS_117004; V2LHS_117217) were purchased from Dharmacon (Horizon Discovery; Waterbeach, United Kingdom). A nonsense scrambled shRNA sequence was used as control. Lentiviral particles were generated by transfection of HEK 293 cells (Cat.# ACC635; DSMZ, Braunschweig, Germany) with Pmd2 (AddGene, Plasmid #12259), lentiviral envelope plasmid psPAX2 (AddGene, Plasmid #12260) and the plasmid containing scramble or against *APOLD1* gene sequences. Lentiviral particles were generated by transfection of HEK 293 cells with the respective plasmid and pLenti-C-mGFP-P2A-Puro Lentiviral Gene Expression Vector (Cat. #PS100093, Origene). Lipofectamine 2000 (Cat.# 11668030; Thermo Fisher Scientific) was used for transfection. Viral particles were collected at least 48 hours after incubation. Cells were transduced 24 hours in the presence of 8 μg/ml polybrene and re-fed with fresh medium the next day. Apold1-KD HUVECs were used in functional assays at least 5 days post-transduction.

### Immunohistochemistry and histology

Muscle samples were harvested and embedded in Tissue-Tek and frozen in liquid N2-cooled isopentane. Frozen muscle cross sections (7-10 μm) were made using a cryostat (Leica CM 1950) and collected on Superfrost Ultra Plus slides (Thermo Fischer Scientific). Cross sections were permeabilized in PBS with 0.5% Triton X-100 and blocked for 1 h at room temperature in PBS with 0.05% Tween-20, 1% BSA (BS). Primary antibody incubations were performed at 4 °C overnight. Slides were subsequently washed in PBS and incubated for 1 hour in blocking buffer with the appropriate secondary antibodies. Nuclei were stained with Hoechst. The following primary and secondary antibodies were used: anti-CD31 (PECAM1) (1:250, AF3628, R&D Systems), anti-α smooth muscle actin Cy3™ conjugated (αSMA) (1:500, C6198, Sigma-Aldrich), isolectin B4, (IB4, 1:500, I32450, Thermo Fisher Scientific), Alexa Fluor 594 donkey anti-goat IgG (H+L) (1:250, A-11058, Thermo Fisher Scientific), Alexa Fluor 568 donkey anti-rat IgG (H+L) (1:250, ab175475, abcam), Alexa Fluor 350 donkey anti-sheep IgG (H+L) (1:250, A-21097, Thermo Fisher Scientific), Alexa Fluor 488 donkey anti-rabbit IgG (H+L) (1:250 dilution, A-32790, Thermo Fisher Scientific), Alexa Fluor 488 donkey anti-goat IgG (H+L) (1:250, A-32814, Thermo Fisher Scientific), Alexa Fluor 350 donkey anti-rabbit IgG (H+L) (1:250 dilution, A-10039, Thermo Fisher Scientific).

Fiber type staining combined with isolectin B4 staining was performed in two steps: First, fiber type staining was performed as described (Masschelein et al., 2020). In short, sections were dried and washed for 5 min in PBS supplemented with 0.05% Triton-X-100 (PBST) and subsequently blocked for 60 min in PBST + 10% goat serum (16210064, Thermo Fisher Scientific). Afterwards a primary antibody cocktail diluted in PBST + 10% goat serum was applied for 120 min against MyHC-I (1:50 dilution, BA-F8 from hybridoma, Iowa City, IA, USA), MyHC-IIa (1:200 dilution, SC-71 from hybridoma) and MyHC-IIb (1:100 dilution, BF-F3 from hybridoma). After washing 3 times for 5 min, a secondary antibody cocktail, diluted in PBST + 10% goat serum, was applied for 60 min. Secondary antibodies were: Alexa Fluor 488 goat anti-mouse IgG2B (1:250 dilution, Thermo Fisher Scientific), Alexa Fluor 350 goat anti-mouse IgG1 (1:250 dilution, A-21120, Thermo Fisher Scientific), Alexa Fluor 568 goat anti-mouse IgM (1:250 dilution, A-21043, Thermo Fisher Scientific). After a 3 x 5 min wash, muscle sections were briefly fixed in 2% PFA for 5 min, and washed with PBS. For additional IB4 vascular staining, muscle sections were incubated overnight at 4 °C with with Isolectin GS-IB4 Alexa Fluor™ 647 (IB4, 1:500, I32450, Thermo Fisher Scientific) diluted in blocking buffer 0.1% Triton X-100. For EdU detection combined with ERG and/or PECAM1/IB4, EdU was first visualized using the EdU Click-iT™ Cell Reaction Buffer Kit (C10269, Thermo Fisher Scientific) according to manufacturer’s instructions, and subsequently incubated for 1 h in blocking buffer (PBS with 1% BSA) at room temperature (RT). Thereafter, sections were incubated overnight at 4 °C with Isolectin GS-IB4 Alexa Fluor™ 568 (1:500, I21412, Thermo Fischer Scientific), goat anti-Mouse/Rat CD31 (PECAM1) antibody (1:250, 3628, R&D Systems) and ERG antibody (#97249, Cell Signaling, 1:200) diluted in blocking buffer with 0.1% Triton X-100.

Images were taken with a Zeiss Axio observer Z.1 or an Olympus confocal microscope (FV1200). All images were captured at the same exposure time for one experiment. Composite images were stitched together using the tiles module in the ZEN 2011 imaging software (Zeiss). Fiber type composition, vascular density (% IB4/PECAM1+ area) and ERG numbers were quantified using ImageJ software.

### *In vitro* analysis of EC function

#### Proliferation

Cultured primary mouse mECs (no longer than 8 days, P1) or HUVECs were incubated in growth medium containing 10 μM 5-ethynyl-2’-deoxyuridine (EdU) for 15 hours. As a measure of proliferation, incorporation EdU was assessed using the Click-iT™ Cell Reaction Buffer Kit (C10269, Thermo Fisher Scientific), according to the manufacturer’s instructions. Briefly, after EdU incorporation, cells were fixed with 4% paraformaldehyde for 10 min and permeabilized for 20 min in 0.5% Triton X-100 with 3% BSA in PBS, followed by reaction with the Click-iT reaction cocktail for 45 min in dark at room temperature. Thereafter, cells were washed briefly and counterstained with Hoechst (#62249, Thermo Fisher Scientific,1:2000) and antibodies against CD31 (PECAM1) (AF3628, R&D Systems, 1:250) or ERG (#97249, Cell Signaling, 1:250). Cells were imaged using a Zeiss Axio Observer.Z1 fluorescence microscope (Carl Zeiss, Oberkochen, Germany). EdU^+^ cells and ERG^+^ ECs were counted in at least 5 random fields and the percentage of EdU^+^ cells in ECs was calculated.

#### Scratch wound assay

A scratch wound was applied on confluent EC monolayers (pre-treated with 1 µg/ml mitomycin C for 24 hrs where indicated) using a 200 µl tip. After scratch wounding (T0) and photography using a Leica DMI6000 B inverted microscope (Leica Microsystems, Mannheim, Germany), the cultures were further incubated in growth medium and fixed with 4% PFA 24 hours after first scratch (T24). Cells were photographed again (T24) and gap areas at both time points were measured using the Fiji software package (https://fiji.sc) to calculate the percentage of wound closure using the following expression: (1 – (T24_gap area_/T0_gap area_)) x 100.

#### Spheroid capillary sprouting assay

Spheroids were prepared as previously described (De Bock et al., 2013) with minor modifications. Briefly, spheroids containing 1000 HUVECs, or mECs per 25 µl droplet were plated overnight as hanging drops in a 20% methylcellulose (9004-67-5, Sigma-Aldrich) in EGM2 mixture. The next day, spheroids were collected in 10% FBS in PBS, concentrated using several centrifugation steps and embedded in a Fibrinogen gel (5 mg/ml fibrinogen (F8630, Sigma-Aldrich) dissolved in EGM2 plus 1U/ml thrombin (T4648, Sigma-Aldrich). To assess tip cell competition, cells were mixed at the indicated ratio. Growth medium (with or without MitomycinC) were pipetted on top of the gel to induce sprouting. 24 hours later, spheroids were fixed with 4% PFA at room temperature and photographed using a Leica DM IL LED microscope (Leica Microsystems GmbH, Wetzlar, Germany). <u>Contact inhibition</u>: HUVECs were seeded in 50% EGM2 / 50% full M199 medium at a density of 15,000 cells/cm^2^ and were further cultured for 3 days until contact inhibition was reached. To generate the corresponding proliferative control, contact inhibited cells were trypsinized and cultured for 24 hr to re-initiate proliferation.

#### VEGF-A stimulation

HUVECs were starved with DMEM medium without any growth factors for 4 hrs. One group was treated with VEGF (Cat.# 450-32; PeproTech; London, United Kingdom)(end conc.: 30 ng/ml) for two hours, while the other group was starved for another two hours. RNA was extracted as described above.

#### Assessment of autophagy

Cultured primary muscle endothelial cells were incubated in either growth medium or medium lacking amino acids for 16 hrs. Cells were collected and lysed with [50 mM Tris–HCl pH 7.0, 270 mM sucrose, 5 mM EGTA, 1 mM EDTA, 1 mM sodium orthovanadate, 50 mM glycerophosphate, 5 mM sodium pyrophosphate, 50 mM sodium fluoride, 1 mM DTT, 0.1% Triton-X 100 and a complete protease inhibitor tablet (C755C25, Roche Applied Science)]. Lysates were centrifuged at 10,000 g for 10 min at 4 °C. Supernatant was collected and protein concentration was measured using the DC protein assay kit (5000116, Bio-rad). Total protein (5-10 ug) was loaded in 15 well pre-casted gradient gel (456-8086, Bio-Rad). After electrophoresis, a picture of the gel was taken under UV-light to determine protein loading using stain-free technology. Proteins were transferred onto a PVDF membrane (Bio-rad, 170-4156) with a semi-dry system. Membrane was blocked for 1h at room temperature with 5% BSA in 0.1% TBS tween. Membranes were incubated in primary antibodies (1:500, NB100-2220, Novus Biologicals; 1:500, ab109012, Abcam,) overnight. Anti-rabbit IgG HRP-linked secondary antibody (1:500, 70748, Cell Signaling Technology) was used for chemiluminescent detection of proteins. Membranes were scanned with a Chemidoc imaging system (Bio-rad) and quantified using Image Lab software (Bio-rad).

### Flow Cytometry

For endothelial cell analysis in muscles, muscles were dissected, separated and enzymatically digested as described above, cells were incubated in dark for 30 min with anti-mouse CD31 (PECAM1) PE antibody (553373, BD Biosciences) and anti-mouse CD45 PerCP antibody (557235 BD Biosciences) (1:400 diluted in FACS buffer (1xPBS+1% FBS)). Cells were washed with FACS buffer before loading. For EdU proliferation experiments in B16-F10 melanoma, tumors were dissected after seven hours labeling with EdU (i.p. 5 mg/ml in saline, 10μg per gram body weight), then the dissociated cells were briefly fixed with 2% PFA and processed with the click-iT plus EdU Alexa Fluor® 647 Flow Cytometry Assay Kit (C10634, ThermoFischer Scientific) according to the manufacturer’s instructions. Subsequently, they were incubated in the dark for 30 minutes with CD31 (PECAM1) PE antibody (553373, BD Biosciences) and Alexa Fluor® 488 anti-mouse CD45 Antibody (1:400, 103122, BioLegend). Cells were analyzed using SONY SH800S cell sorter. Data were analyzed using FlowJo 10 software (Tree Star).

### Quantification and statistical analysis

The images presented in the manuscript are representative of the data (quantification of image is approximately the group average) and the image/staining quality. All data represent mean ± SEM. GraphPhad Prism software (version 8.0.0) was used for statistical analyses. Investigators were always blinded to group allocation. Unless otherwise indicated, when comparing two group means, Student’s t-test was used in an unpaired two-tailed fashion. For more than two groups, one-way ANOVA with Tukey’s multiple comparisons test was used and for experimental set-ups with a second variable, two-way ANOVA with Sidak’s multiple comparisons test and two-way ANOVA with Tukey’s multiple comparison test was used. The statistical method used for each experiment is indicated in each figure legend. Asterisks in figure legends denote statistical significance. No experiment-wide multiple test correction was applied. P>0.05 is considered non-significant (n.s.). P<0.05 is considered significant (*).

---

**Fig. S1.**
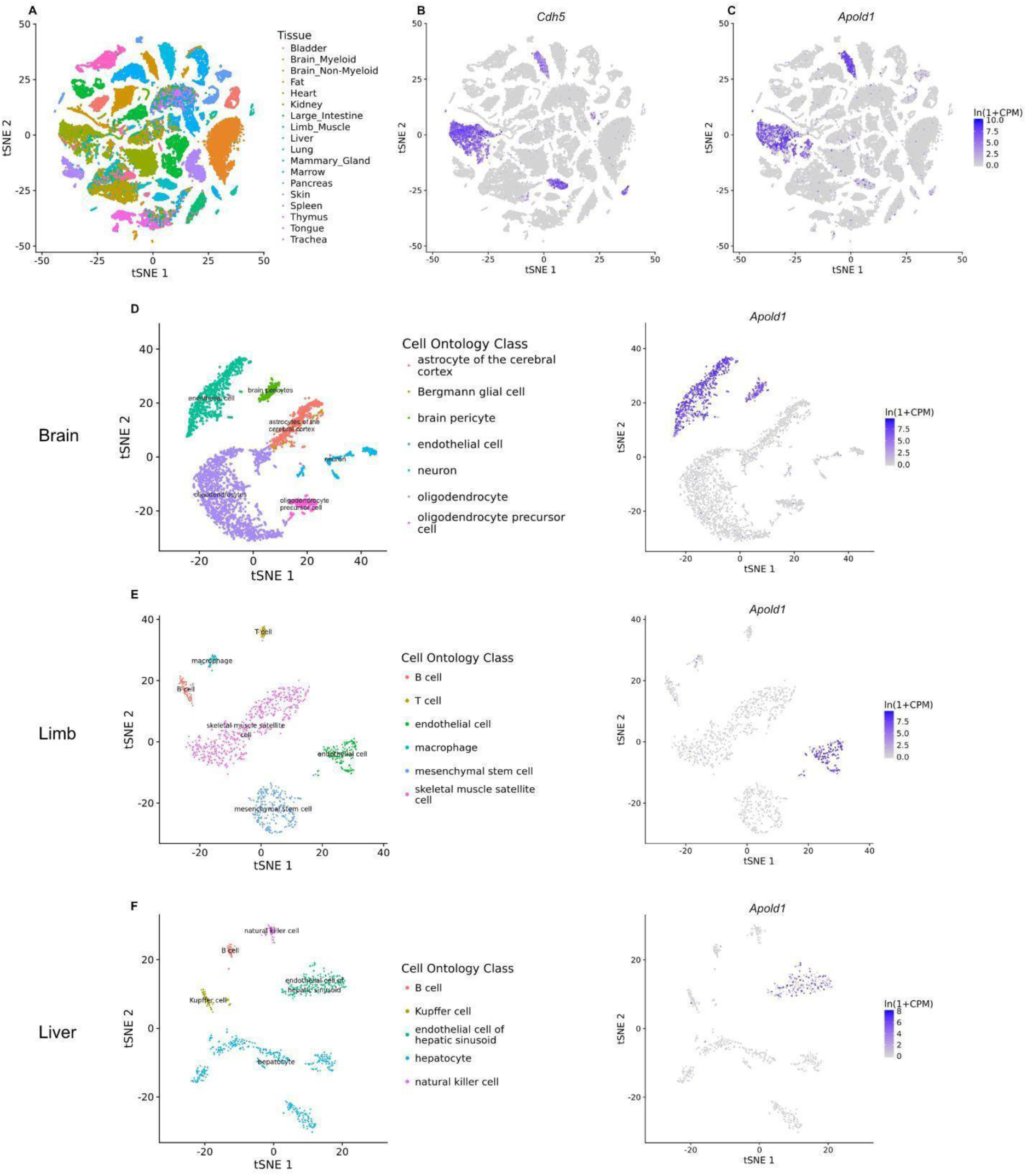
Across multiple organ systems in the mouse, *Apold1* expression is largely restricted to endothelial cells. (A) Analysis of the ’Tabula Muris’ compendium of FACS sorted data across organs [34]. (B) Endothelial cells identified using *Cdh5* expression co-localize with (C) *Apold1* expression. (D) Across organs, endothelial cells in various tissues are highly enrichted in *Apold1*. Restricting the ’Tabula Muris’ dataset to FACS sorted cells, the *Apold1* enrichment in ECs is apparent in various highly vascularized tissues, including (E) the brain, (F) muscles, and (G) the liver.

**Fig. S2.**
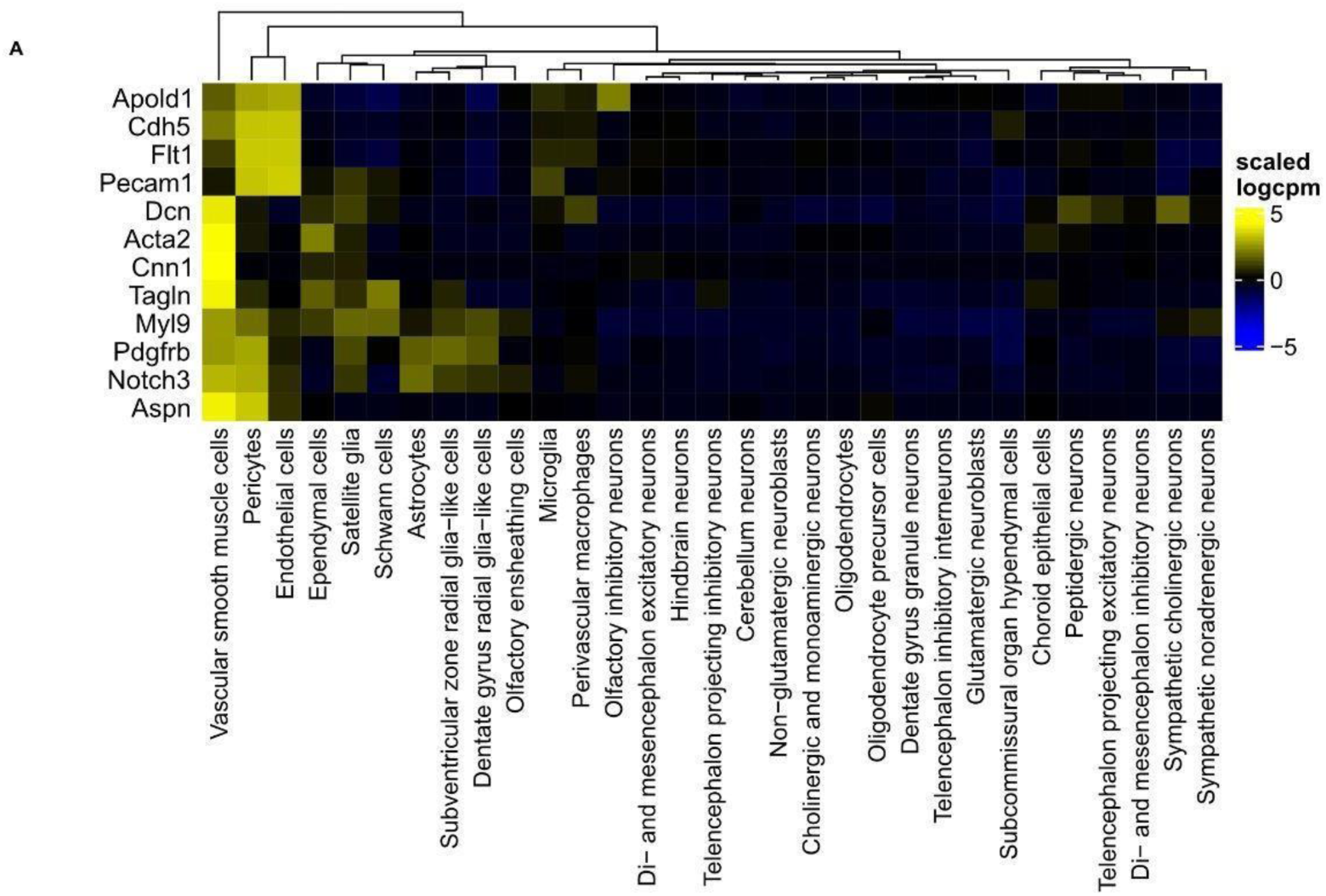
A*p*old1 expression in the Zeisel et al 2018 study, an extensive single-cell characterization of the mouse nervous system [38].

**Fig. S3.**
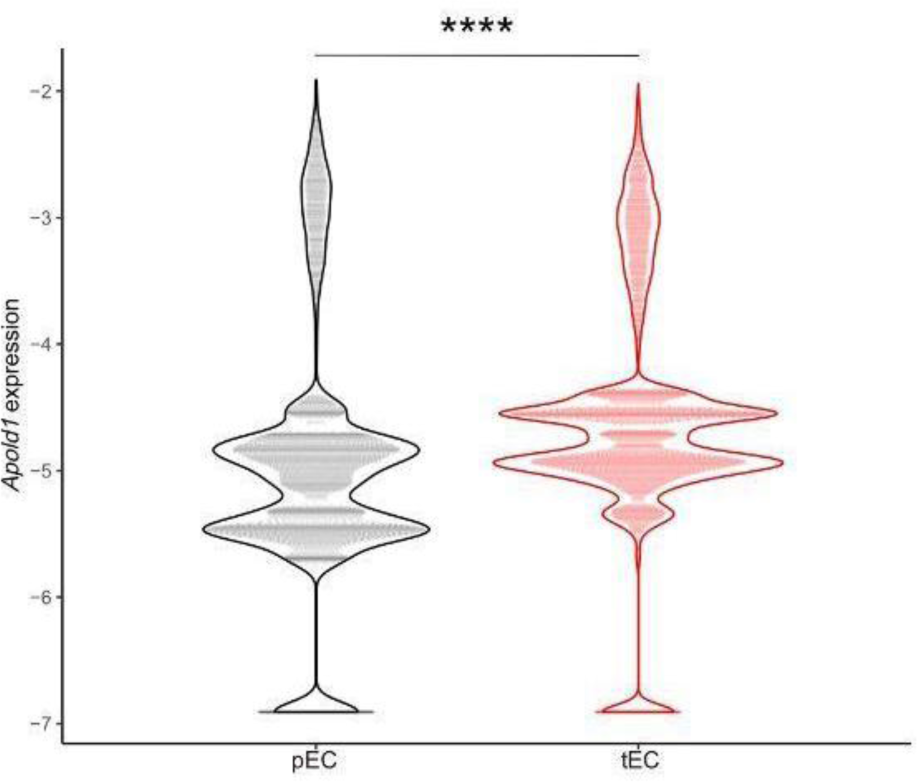
High *Apold1* expression in tumor ECs and specifically in tip cells. scRNA-seq data reanalyzed [52]. Freshly isolated human tumor ECs (TECs) were analyzed in direct comparison to peritumoral pulmonary non-tumor ECs (PNECs) from the same patient. Data come from 1 large cell carcinoma, 4 squamous cell carcinomas, and 3 adenocarcinoma treatment-naive patients. (A) *Apold1* expression in TECs versus NECs (pEC mean= -4.9; tEC mean = -4.58; **** p < 0.0001).

**Fig. S4.**
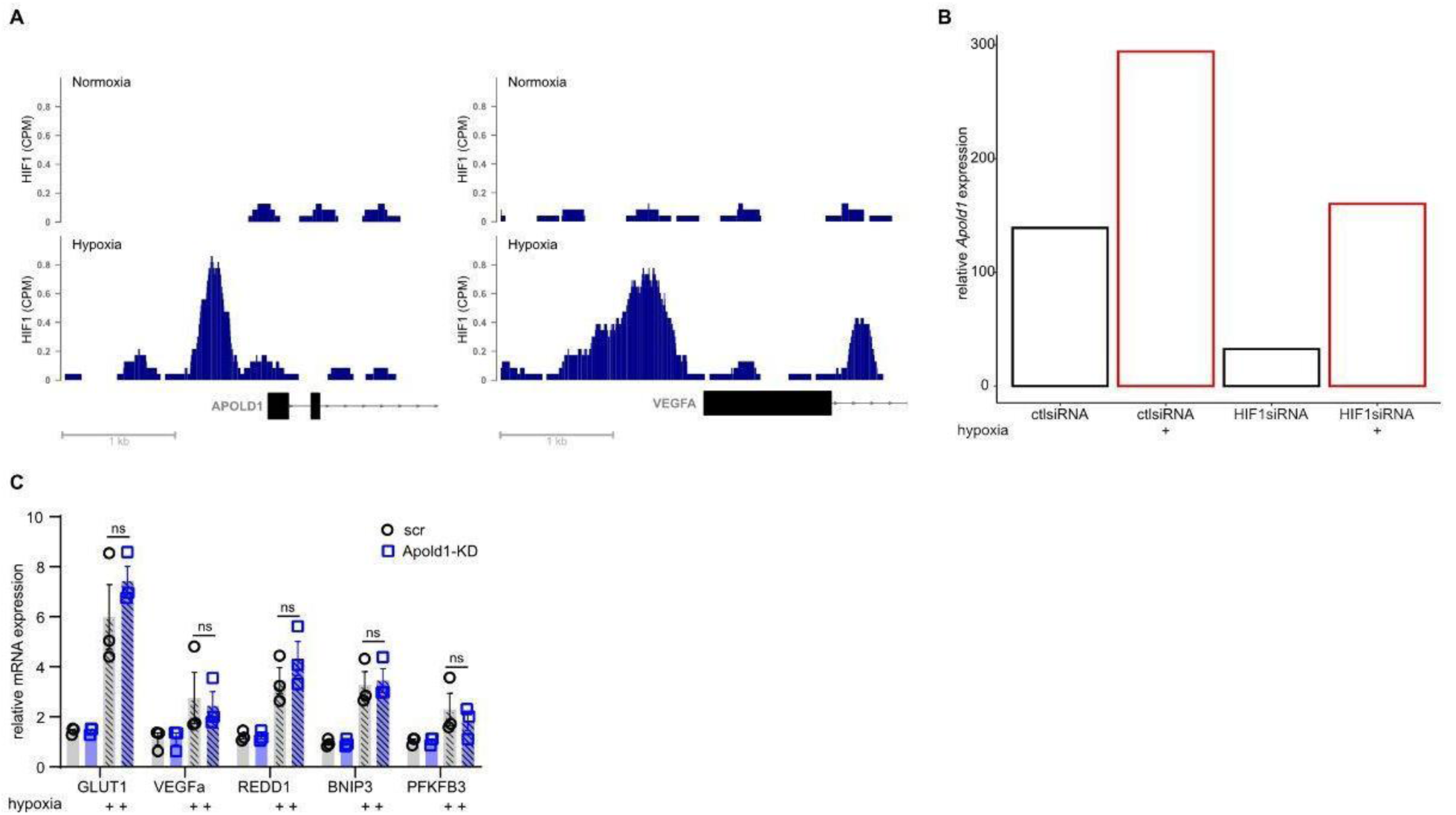
HIF1 knockdown prevents the hypoxia-induced increase in Apold1 in HUVECs. (A) Hypoxia dependent HIF binding sites to the promoter region of Apold1 and VEGF reanalyzed from a HIF1-ChIP-seq data-set [55]. (B) *Apold1* expression in HUVECS in response to Hif1a-knockdown under normoxia and hypoxia reanalyzed [55]. (C) RT-qPCR results for canonical hypoxia-responsive mRNAs in scr and Apold1-KD under normoxia (21% O2) and hypoxia (0.1% O2) for 16 hrs (n (scr/Apold1-KD) = 3/3). Student’s t-test in C. The data shown are mean ± SEM.

**Fig. S5.**
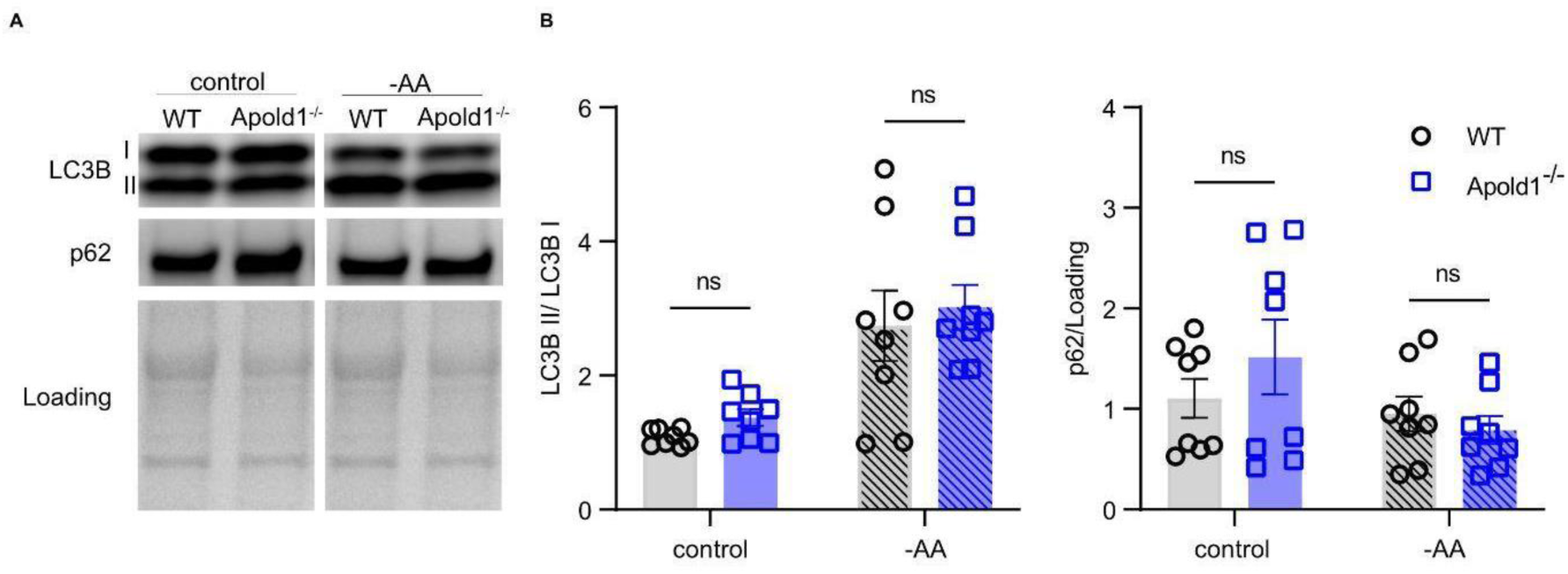
Apold1 deficiency is not affecting autophagic markers. (A) Representative images of western blot analysis of autophagic markers in control and amino acid (-AA) deprivation conditions in cultured WT and *Apold1^-/-^* mECs. (B) Quantification of LC3B ratio and p62 in control and amino acid deprivation (n (WT/*Apold1^-/-^*) = 8/8). Student’s t-test in B. The data shown are mean ± SEM.

## Notes

### Competing Interest Statement

The authors have declared no competing interest.

